# Desert Dingo (*Canis lupus dingo*) genome provides insights into their role in the Australian ecosystem

**DOI:** 10.1101/2020.11.15.384057

**Authors:** Sonu Yadav, Olga Dudchenko, Meera Esvaran, Benjamin D. Rosen, Matt A. Field, Ksenia Skvortsova, Richard J. Edwards, Shyam Gopalakrishnan, Jens Keilwagen, Blake J. Cochran, Bikash Manandhar, Martin Bucknall, Sonia Bustamante, Jacob Agerbo Rasmussen, Richard G. Melvin, Arina Omer, Zane Colaric, Eva K. F. Chan, Andre E. Minoche, Timothy P.L. Smith, M. Thomas P. Gilbert, Ozren Bogdanovic, Robert A. Zammit, Torsten Thomas, Erez L. Aiden, J. William O. Ballard

## Abstract

The dingo is Australia’s iconic top-order predator and arrived on the continent between 5,000-8,000 years ago. To provide an unbiased insight into its evolutionary affiliations and biological interactions, we coupled long-read DNA sequencing with a multiplatform scaffolding approach to produce an *ab initio* genome assembly of the desert dingo (85X coverage) we call CanLup_DDS. We compared this genome to the Boxer (CanFam3.1) and German Shepherd dog (CanFam_GSD) assemblies and characterized lineage-specific and shared genetic variation ranging from single– to megabase pair–sized variants. We identified 21,483 dingo-specific and 16,595 domestic dog-specific homozygous structural variants mediating genic and putative regulatory changes. Comparisons between the dingo and domestic dog builds detected unique inversions on Chromosome 16, structural variations in genes linked with starch metabolism, and seven differentially methylated genes. To experimentally assess genomic differences 17 dingoes and 15 German Shepherd dogs were fed parallel diets for 14 days. In dingoes, low *AMY2B* copy number and serum amylase levels are linked with high cholesterol and LDL levels. Gut microbiome analyses revealed enrichment of the family *Clostridiaceae*, which can utilize complex resistant starch, while scat metabolome studies identified high phenylethyl alcohol concentrations that we posit are linked with territory marking. Our study provides compelling genomic, microbiome, and metabolomic links showing the dingo has distinct physiology from domestic breed dogs with a unique role in the ecosystem.

## Main

Australia has the worst mammalian extinction rate of any country in the world and the catastrophic bushfires of 2019-20 have fast tracked multiple species towards extinction. Concomitant with public education a strategic priority must be to restore ecosystem balance. One approach to restoring ecosystems and to conferring resilience against globally threatening processes is to develop our understanding of the functionality of predators^1^. Dingoes have been the Australia’s apex predator since their arrival 5,000-8,000 years ago^2,3^. They show a unique suite of behavioural traits including scent-marking for social communication, territory defence and to synchronise reproduction. Historically, they fed on a marsupials and reptiles. In native ecosystems, they tend to consume the most prevalent species^4^. In disturbed environments dingoes eat prey of increasing body size as aridity increases^5^. This opportunistic hunting has brought the dingo into conflict with pastoralists and feral dogs.

To resolve the debate around the ecological role of dingoes in the Australian ecosystem it is crucial to identify the structural and functional genetic differences that distinguish them from feralised domestic dogs. To date, genomic studies have been based on mapping re-sequenced genomes to the domestic dog reference genome^6–9^. The alignment of re-sequenced data to a single reference genome underestimates species-specific variation, yet computational analyses have established the dingo genome harbours multiple positively selected genes related to metabolism ^6,10,11^. Further, dingoes have retained the ancestral pancreatic amylase *AMY2B* copy number (n=2) with one or more copy number expansions in domestic dogs^10^. We explore the genomic divergence between a desert dingo and two domestic dog breeds and experimentally consider whether differences in the biochemistry, physiology and digestive gut microbiome influence organismal functions and ecological roles.

We assemble the genome of a wild-found dingo named “Sandy” (Fig. 1a) and compare it with the Boxer (CanFam3.1)^12^ and German Shepherd Dog (GSD) (CanFam_GSD)^13^. The boxer is a highly derived, brachycephalic breed with a mesocephalic head shape ^13^. GSDs are intermediate in the currently accepted modern domestic dog phylogeny^14^, are morphologically similar to dingo with medium body size and are common on farms. GSD crossbreds are also common feral dogs. We conducted structural variation analyses, genomic selection scans, and DNA methylation studies to identify dingo genomic features. To examine the influence of the distinct evolutionary histories on organismal physiology, we experimentally compared dingo and GSD serum, gut microbiome, and scat metabolites. Our study uncovers compelling evidence to suggest the Australian dingo has a unique role in the ecosystem that is mediated by its evolutionary history and ancient divergence from domestic dog breeds.

**Fig. 1:**
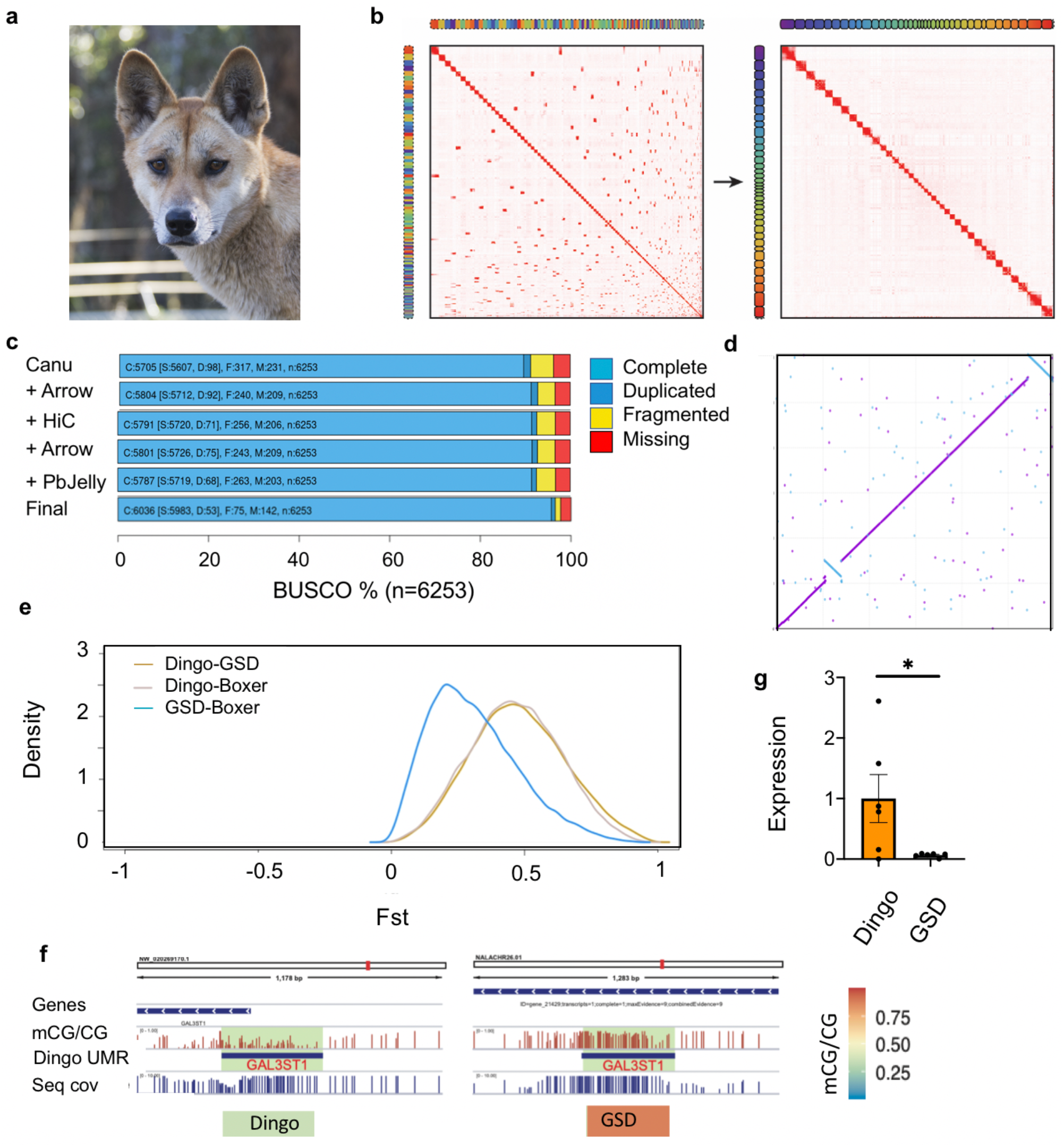
Sandy the Desert dingo and her genome. **a**, Sandy as a 3-year-old. She was found as a 4-week old puppy in a remote region of South Australia in 2014. Subsequent genetic testing showed she was a pure desert dingo. **b**, Contact matrices generated by aligning the Hi-C data set to the genome assembly before Hi-C scaffolding (left), and after Hi-C scaffolding (right). Interactive contact matrices are available on www.dnazoo.org/assemblies. **c**, BUSCO v3 completeness scores for different stages of the genome assembly (C: complete, S: single, D: duplicated, F: fragmented, M: missing). **d**, Synteny plot for chromosome 16 CanFamv3.1 (x-axis) vs dingo (y-axis). The dingo assembly contains two large inversions relative to CanFamv3.1. **e**, Fst distribution for Dingo-GSD, Dingo-Boxer and GSD-Boxer (dingo *n* = 10, GSD *n* = 20 and Boxer *n* = 14). **f**, DNA methylation differences at transcription start sites (TSS) proximal regulatory regions (UMRs) in *GAL3ST1* between the dingo and GSD. Heatmap showing DNA methylation levels at TSS-associated UMRs, differentially methylated between dingo and GSD. IGV browser tracks depicting DNA methylation differences at UMRs between the dingo and GSD. **g**, Significant difference in expression of *GAL3ST1* between dingo and GSD (t_(10)_ =2.361, P= 0.03, dingo *n* = 6, GSD *n*= 6). Mean SE is shown on the plot. * shows P<0.05.

### Genome assembly, annotation, and comparative analyses

Genomic DNA was extracted from a pure female dingo found in the Strzelecki Desert in South Australia. The genome was assembled using a combination of long-read sequencing approaches with Hi-C scaffolding (Fig. 1b; Extended Data Fig. 1). The assembly has a size of 2.35 Gb, consists of 159 scaffolds with a contig and scaffold N50 length of 64.3 Mb (contig L50=20, scaffold L50=14) and 33.7 kb of gap sequence (Supplementary Information 1). The full-length chromosome scaffolds in the assembly accounted for 99.46 % of the genome. In total 93.0 % of the conserved single-copy genes were complete. Compiling BUSCO results across all assembly stages reveals at least 6,036 conserved genes (96.5 %) are present and complete in the assembly, with only 142 genes (2.27 %) not found (Fig 1c, Supplementary Table 1.1, 1.2). BUSCO analysis of the longest isoform per annotated gene increased this number to 6,174 (98.7%) complete with only 18 (0.3%) missing (Supplementary Table 1.1). KAT kmer analysis showed no sign of missing data nor large duplications (Extended Data Fig. 2).

Considering the major chromosome alignments, the dingo assembly covers 99.16% of the CanFam3.1 assembly compared to 99.31% of the CanFam_GSD assembly. Conversely, 99.03% of the dingo aligns with CanFam3.1, while 98.54% of the CanFam_GSD assembly aligns to CanFam3.1. These differences are largely attributable to ~38 Mb of extra sequence in CanFam_GSD relative to CanFam3.1 compared to only ~1Mb of extra sequence in the dingo assembly. Synteny plots were generated for each chromosome and overall there were limited large-scale genomic rearrangements. Chromosome 16 however contained two large inversions in the dingo compared to CanFam3.1 (Fig. 1d) and one large inversion CanFam_GSD vs CanFam3.1 (Extended Data Fig. 3) indicating differential evolutionary signatures in dingoes compared to other canid lineages.

Several approaches were employed to assess the level of variation in the dingo genome (Supplementary Information 1.9). Small-scale variations (SV), generally <50 bp, were detected in both the dingo assembly and CanFam_GSD relative to CanFam3.1. Overall, a total of 4.5 k SNPs were called in dingo compared to 3.6 k SNPs in CanFam_GSD, representing 22% more SNP calls in dingo. Additionally, there were 6.2 k small indels detected in the dingo compared to 5.1 k small indels in CanFam_GSD representing 21% more small indel calls.

Relative to CanFam3.1, a total of 75.8 k SVs were detected using Nanopore reads and 116.2 k SVs were detected using PacBio reads. Fewer SVs were detected overall relative to CanFam_GSD with a total of 63.8 k SVs detected using Nanopore reads and 99.1 k SVs detected using PacBio reads. To account for higher SV false-positive rates, a more conservative list of SVs was generated consisting of the intersection of PacBio and Nanopore calls using a consensus approach^15^. This resulted in 73.5 k CanFam SVs and 62.4 k CanFam_GSD SVs, of which over 99% are either insertions or deletions. To prioritise structural variants for further investigation, SVs were overlapped to existing CanFam3.1 gene annotations and dingo gene annotations generated with GeMoMa (version 1.6.2beta)^16^. With the CanFam SVs, 29,688 were found to overlap protein-coding genes compared to 26,760 for CanFam_GSD SVs. These SVs were then filtered for homozygous events yielding 24,515 CanFam3.1 SVs (representing 8571 unique genes) compared to 21,961 CanFam_GSD SVs (representing 7,650 unique genes). The remaining deletions (insertions) represent 13.94 (2.97) Mb of total deleted sequence relative to CanFam3.1 and 5.03 (1.79) Mb relative to CanFam_GSD.

The prioritised SV’s were next examined for overlap to specific genes of interest. We examined all structural variant calls overlapping *AMY2B* as variation in copy number has been linked to starch diet adaptations^17^. A single SV was detected, a heterozygous 203 bp deletion detected in the PacBio dingo reads relative to CanFam_GSD, which contains 7-8 copies of *AMY2B*^13^. This *AMY2B* SV indicates the possibility of diversification of the gene involved in starch digestion between dingoes and other canids. A broader analysis was performed overlapping the regions previously identified as important in dog domestication^18^. In total 132 SVs were identified that overlapped these regions containing 44 unique genes (Supplementary Table 1.3), including *MGAM*, which is also involved in starch metabolic and catabolic processes.

To quantify the genetic differentiation and signatures of selection across the genome between dingoes and two domestic dog breeds, we computed the pairwise Fst between the dingo, Boxer, and GSD (Supplementary Information 1.10). We did not include additional breeds because alignments of short read sequences to distinct *de novo* assembles can cause bias^19^. Fst distribution of dingo-GSD and dingo-Boxer differed from GSD-Boxer (Fig. 1e). As expected, selection scan indicated higher genetic differentiation in the dingo-GSD, dingo-Boxer than GSD-Boxer for *AMY2B* and *MGAM* (Extended Data Fig. 4).

Next, we compared the DNA methylation of Sandy the dingo and Nala the GSD^13^. DNA methylation status of the transcription start sites (TSS) associated regulatory regions may serve as a proxy for the activity of the corresponding gene. The highly methylated gene promoters are often indicative of a transcriptionally repressed state while unmethylated gene promoters indicate a transcriptionally permissive state. In our study, five unmethylated regions with genes: *GAL3ST1, NAP1L5, FAM83F*, *MAB21L1*, and *UPK3A* showed reduced DNA methylation in the dingo translating to their higher expression levels (Fig. 1f, 1g) and two UMRs *LIME1* and *GGT5* showed hypermethylation in dingo (Extended Data Fig. 5). Of these, *GAL3ST1* is associated with galactose metabolism by catalysing sulfation of galactose^20^. Dingo and dingo-dog hybrids differ in their galactose metabolism likely linked with differences in *AMY2B* copy number^21^.

Assembly, annotation and comparative analyses of the desert dingo genome shows that it has forked from that of the Boxer and GSD. Likely this is due to the ancient divergence of the dingo from the domestic breeds, recovery of genetic variation since dingoes colonised Australia 5,000-8,000 years ago and selection for feeding on marsupials with low fat and high protein meats. In the next section, we conduct a dietary manipulation study to link the dingo and GSD genomes with organismal biology to gain insight into the roles of dingoes and feral dogs in the ecosystem (Supplementary Table 2.1).

### Biochemical, physiological, and microbiome differences between dingoes and GSDs

Prior to the dietary manipulation study, we minimised variation in the gut flora by treating canids with a broad-spectrum antibiotic and then supplementing their diets with a probiotic. In parallel, 17 dingoes and 15 GSDs were fed a constant diet for 10d and then the proportion of rice was increased to 75% over the next 4d (Supplementary data 2.2). As expected, ddPCR analysis showed *AMY2B* copy number and serum amylase levels were lower in dingoes than GSDs (Fig 2a, b). Unexpectedly, total cholesterol was significantly higher in the dingoes as compared to GSDs (Fig 2c). Low-density lipoprotein (LDL) cholesterol was elevated in dingoes (Fig 2d), but there were no obvious differences in high-density lipoprotein cholesterol levels or in lipase or triglycerides (Extended Data Fig. 6). Elevated cholesterol and LDL levels are protective against infection^22^, suggesting dingoes have an elevated immune response in comparison to GSDs^23^.

**Fig. 2:**
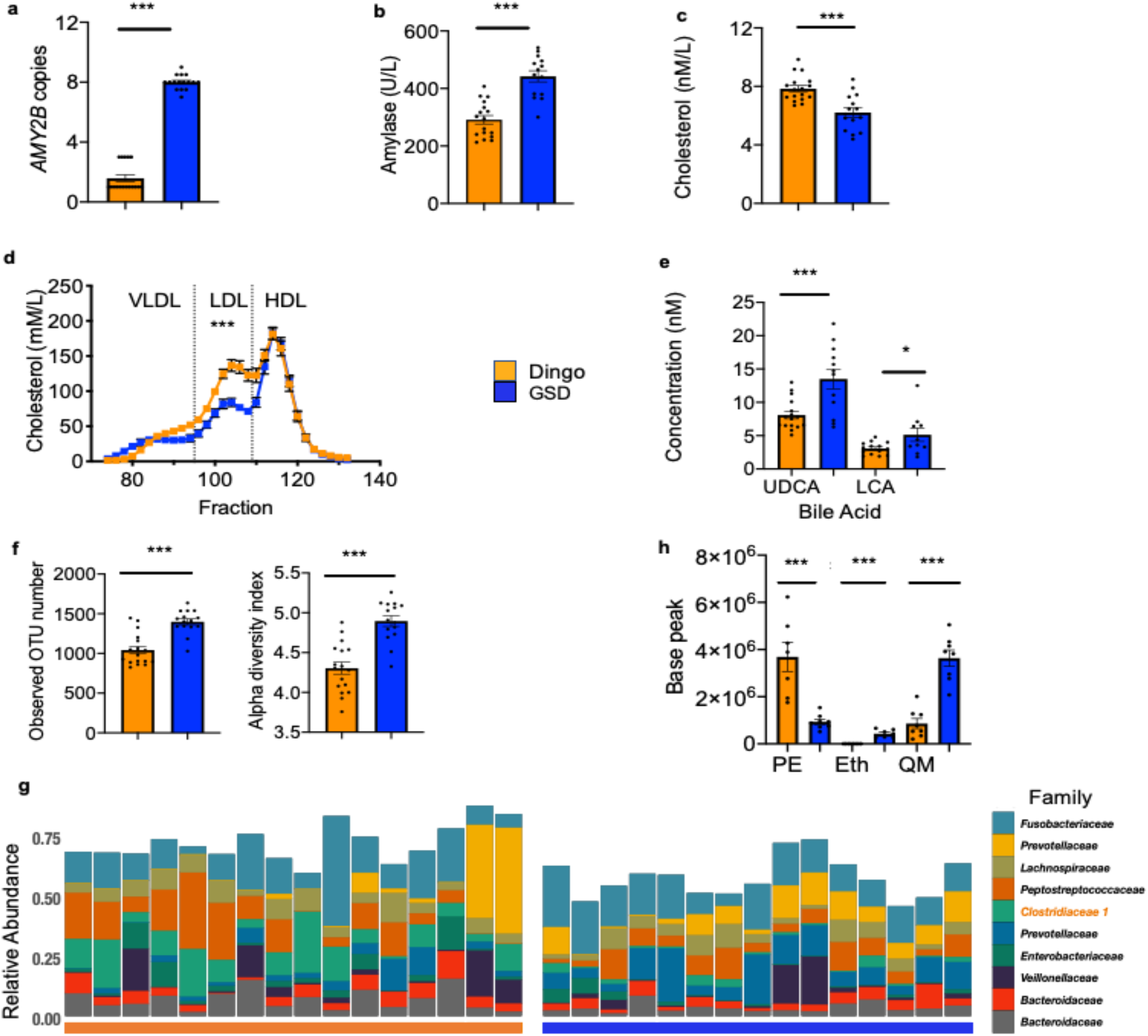
Biochemical and physiological differences between dingoes and German Shepherd Dogs (GSD). **a**, Amylase. *AMY2B* copy number is lower in dingoes than GSDs (t_31_ =24.42, P<0.0001) with fewer copies in dingoes (mean=1.58 ± 0.22) than GSDs (mean=8 ± 0.12; dingo *n* = 17, GSD *n* = 16). **b**, Serum amylase levels are lower in the dingo compared to the GSD (t_29_ =6.25, P<0.0001; dingo *n* = 17, GSD *n* = 14). **c**, Total cholesterol is significantly higher in the dingoes as compared to GSDs (t_30_ =4.36, P=0.0001; dingo *n* = 17, GSD *n* = 15). **d**, LDL-C is elevated 2.2-fold in dingoes (t_10_=4.64, P< 0.001; dingo *n* = 6, GSD *n* = 6) but no obvious difference in HDL-C levels. Individual points are within symbol size. **e**, Two secondary bile acids Ursodeoxycholic acid (UDCA) (t_26_=3.732, P<0.001; dingo *n* = 16, GSD *n* = 12), and Lithocholic acid (LCA) (t_22_ =2.314, P= 0.030; dingo *n* = 14, GSD *n* = 10) are significantly lower in dingoes. **f**, Microbial diversity: LHS Microbial richness (Wilcoxon Rank Sum test P.adj =0.00003; dingo *n* = 17, GSD *n* = 15). RHS. Shannon’s diversity in the dingo and GSD (Wilcoxon Rank Sum test P.adj =0.00001; dingo *n* = 17, GSD *n* = 15). **g**, Relative abundance of the top 10 most abundant zOTUs at completion of the diet study on the y-axis for the dingoes (*n* = 16) and GSDs (*n* = 15) along the x-axis. *Clostridiaceae 1* is highlighted in the legend as it is elevated in dingoes. **h**, Metabolite differences between dingo and GSD in the scat; PE= Phenylethyl alcohol (t_13_=4.68, P=0.0004; dingo *n* = 7, GSD *n* = 8), Eth= Ethanone, 1-phenyl (t_13_ =7.26, P<0.0001; dingo *n* = 7, GSD *n* = 8),) and QM= Quinoline, 2-methyl (t_14_=6.88, P<0.0001; dingo *n* = 8, GSD *n* = 8). Mean SE is shown on the plot. * shows P<0.05, ** P<0.01, *** P<0.01.

A significant difference in cholesterol levels leads to the prediction that bile acid levels would differ between canids as primary bile acids are synthesized from cholesterol^24^. We observed no significant difference in the concentration of primary bile acids, however, levels of the secondary bile acids ursodeoxycholic acid (UDCA) and lithocholic acid (LCA) were higher in GSDs (Fig 2e; Supplementary Table 2.2). High levels of UDCA and LCA are involved in immune suppression^25^. They also influence the gut microbial community^26^, and may lead to diseases of the gastrointestinal system^24^.

Amylase, cholesterol, and bile acid levels can shape the gut microbiome so we investigated scat microbial communities^26,27^. There was a trend for reduced diversity and richness in the scat microbial community of dingoes on day one of the dietary study (Extended Data Fig. 7a). On day 14, dingoes had markedly reduced microbial richness and diversity (Fig. 2f), with a distinct microbial community structure and composition (Extended Data Fig. 7b). Aligning with our previous observations (Fig. 2c), microbial communities in dingoes show higher metabolic potential for cholesterol and protein metabolism and lower metabolic potential for bile secretion (Extended Data Fig. 7c).

Analysis of the microbiome composition showed that one microbial phylum, 17 families, and 51 genera differed between the canids (Supplementary Table 2.3, 2.4). In dingoes, the family *Clostridiaceae* and the genus *Clostridium sensu stricto 1* were enriched (Fig. 2g). *Clostridium sensu stricto 1* can utilize complex resistant starch^28^ that will not have been broken down by the dingoes due to low amylase activity. In contrast, bacteria of the families *Lactobacillaceae*, *Ruminococcaceae*, and *Prevotellaceae* were depleted in dingoes (Extended Data Fig. 7d, 7e), although two dingoes from Pure Dingo from had high numbers of the latter Family suggesting environmental differences may also be important. These three families are involved in the fermentation and degradation of starch products^29–31^. Linking with our observation that dingoes have high cholesterol (Fig. 2c) the genera *Lactobacillus* and *Eubacterium* were low in dingoes. Specific strains of these taxa have a demonstrated capacity to reduce cholesterol levels^26,32–34^.

We hypothesised that the dingo genome and microbiome will influence scat metabolites and the composition of chemicals involved in territory marking. We found three chemical differences between the two groups. Phenylethyl alcohol (PE) is elevated in dingoes, while ethanone, 1-phenyl quinoline (also known as acetophenone), and 2-methyl (QM) levels are lower (Fig 2h, Supplementary Table 2.5). PE is known to have antibacterial activity, inhibiting the growth of Gram negative bacteria^35^ and elevating levels of *Clostridiaceae*^36^. PE levels are negatively correlated with *Lactobacillaceae*^36^. Acetophenone and 2-methyl have a distinct odour and have previously been shown important chemicals for scent marking in canids^37–39^. Experimental studies are required to test whether the well-established dingo scent-marking behaviour is related to the balance of these three compounds.

## Discussion

Dingoes are a part of the fabric of Australian culture, touching both indigenous groups and more recent immigrants40. They are considered a “*lightning-rod*” of the land as it generates polarised opinions from Aboriginal people, tourism operators, pastoralists, ecologists, conservationists, and evolutionary biologists4. Our comprehensive study underpins the dingoes genomic and ecological distinction from breed dogs by integrating genomic, metabolome, and microbiome analyses. Our genome assembly has high contiguity with few gaps compared to other canine long-read sequencing assemblies (Supplementary Table 1.1). We found unique inversions on Chromosome 16 in the dingo indicating differential evolutionary signatures. Epigenetics analysis indicated seven genes are differentially methylated in the dingo compared to the domestic GSD. Our organismal studies provide insights into the distinct physiology of dingoes as compared to domestic dogs and suggest they have a heightened immune response and a microbial community that, at least partially, compensates for their reduced *AMY2B* copy number.

The importance of dingoes in Australia can be illustrated by comparisons from either side of the Dingo Fence: the world’s largest chain link fence that is designed to keep dingoes out of prime livestock farming country in South-East Australia. Inside the fence, kangaroo populations have skyrocketed, while populations outside the fence are smaller but stable. Excessive kangaroo numbers can overgraze the landscape, compete with livestock and damage vegetation. Further studies of scat metabolites linked to territory marking may prove part of a broad solution to chemically subdivide the landscape and reduce conflict between native animals and commercial farming.

## Methods

Full details of methods can be found in the Supplementary information.

### Genome assembly, annotation, and comparative analyses

The genome was assembled using Pacific Bioscience (PacBio) Single Molecule Real-Time (SMRT) sequencing, Oxford Nanopore (ONT) PromethION sequencing, 10X Genomics Chromium genome sequencing and Hi-C scaffolding (Fig. 1b). Contigs were assembled using SMRT and ONT sequencing ^41^ and then polished ^42^ to minimise error propagation (Supplementary Information 1). To increase the contiguity of the assembly we used the SMRT and ONT reads to fill gaps, which was then followed by a final round of polishing including aligning the 10X Chromium reads to the assembly and Pilon polishing. The resulting chromosome-length genome assembly has been deposited to NCBI (GCA_003254725.2). In addition to the nuclear genome, the mitochondrial genome has been submitted (ID 2385777) and will be linked with the bioproject and biosample.

The CanLup_DDS (Desert Dingo Sandy) and CanFam_GSD assemblies were aligned to CanFam3.1 using MUMmer4^43^ (v4.0.0 beta 2) to assess the overall alignment of the two assemblies. The genome was annotated using the homology-based gene prediction program GeMoMa (GeMoMa, RRID:SCR 017646) v1.6.2beta ^16^ and 9 reference organisms^13^.

Small-scale variation was detected in both the dingo assembly and CanFam_GSD relative to CanFam v3.1 using pairwise MUMmer4^43^ (v4.0.0 beta 2) alignment databases (Supplementary Information 1). To identify large structural differences in the dingo genome, structural variants from both Oxford Nanopore and PacBio sequence data were called relative to CanFam 3.1 and CanFam_GSD.

Genetic differentiation between the dingo and the domestic dog breeds was detected using pairwise Fst on published dingo, GSD, and Boxer genomes (Supplementary Information 1). The short reads were aligned against the dingo de novo reference using the PALEOMIX pipeline^44^. Fst between each pair of populations: dingo-GSD, dingo-boxer, and GSD-boxer was computed using vcftools v0.1.16.

We profiled DNA methylation of the dingo and GSD genomes using MethylC-seq^45^ and identified CpG-rich unmethylated regions (UMRs) overlapping transcription start sites (TSS) in both genomes (Supplementary Information 1). To compare the DNA methylation status of gene promoters between dingo and GSD, we lifted over dingo UMRs to the GSD genome and GSD UMRs to the dingo genome and calculated corresponding DNA methylation. To validate the difference in expression in *GAL3ST1* and *MAB21L1* we performed quantitative reverse transcription PCR RT-qPCR on six dingoes and six GSDs.

### Biochemical, physiological, and microbiome differences between dingoes and GSDs

Before carrying out experiments, diets of the animals were standardised (Supplementary Information 2). Amylase DNA copy number variation was determined using droplet digital PCR (ddPCR) on QX100 ddPCR system (Bio-rad). Amylase, cholesterol, triglycerides, and lipase and were assayed using the Thermo Scientific Konelab Prime 30i at the Veterinary Pathology Diagnostic Services Laboratory (VPDS). Free bile acids in the plasma were quantified using liquid chromatography-tandem mass spectrometry (LCMS/MS) assay.

Simultaneously with the biochemical studies, we sampled scat from the same dingoes and GSD’s on day 1 and 14. DNA was extracted from thawed stool samples (0.3g) using the Qiagen Powersoil kit (cat# 1288-100; Hilden, Germany) according to the manufacturer’s instruction. Library preparation and pair end sequencing was performed (2×300 cycles) on the Illumina MiSeq platform. 16S rRNA gene sequence data were quality filtered and processed for taxonomic assignment and functional predictions (Supplementary Table 2.3)

We examined scat volatile organic compounds (VOC’s) differences using solid-phase microextraction (SPME) gas chromatography-mass spectrometry (GC-MS) (Supplementary Table 2.5).

## Data availability

The complete assembled genome is available at NCBI (ASM325472v2; GenBank assembly accession No. GCA_003254725.2).

## Acknowledgements

Sandy dingo was rescued by Barry and Lyn Eggleton. We wish to thank all those that voted in the PacBio 2017 World’s Most Interesting Genome Competition and to Emily Hatas for running the show. PacBio sequencing was completed by Dave Kudrna at Arizona Genomics Institute. v1.0 of the Genome assembled by Christian Dreischer at Computomics. The 10X and PromethION sequencing was completed at the Garvan Institute, Sydney. Vanessa M. Hayes at the Garvan Institute funded the BioNano data generation used in the v1 Sandy genome assembly. The Hi-C sequencing and chromosome-length assembly were performed by the DNA Zoo Consortium (www.dnazoo.org). For the experimental study, dingoes were made available by Bargo Dingo Sanctuary and Pure Dingo. German Shepherds were kindly supplied by Kingvale and Allendell Kennels. Sam Towarnicki contributed to the biochemical assays and Amy Shaw collected scat. William Donald (UNSW) provided chemical standards for the metabolomics assays. Mass spectrometric results were obtained at the Bioanalytical Mass Spectrometry Facility within the Mark Wainwright Analytical Centre of the University of New South Wales. This work was undertaken using infrastructure provided by NSW Government co-investment in the National Collaborative Research Infrastructure Scheme (NCRIS) subsidised access to this facility is gratefully acknowledged. The project was funded by the Australian Research Council Discovery Project DP150102038. M.T.P.G. and S.G were supported by the ERC (681306 Extinction Genomics) and the Danish National Research Foundation (DNRF143). E.L.A. was supported by an NSF Physics Frontiers Center Award (PHY1427654), the Welch Foundation (Q-1866), a USDA Agriculture and Food Research Initiative Grant (2017-05741), an NIH 4D Nucleome Grant (U01HL130010), and an NIH Encyclopedia of DNA Elements Mapping Center Award (UM1HG009375).

## Contributions

J.W.O.B. coordinated, designed and funded the project. S.Y. compiled the data. R.A.Z. provided the samples. B.D.R. performed the ONT sequencing, genome assembly, and polishing. The DNA Zoo initiative including O.D., A.O., and Z.C. performed and funded the Hi-C experiment. O.D. and E.L.A. conducted the Hi-C analyses. K.S. and O.B. funded and conducted the DNA methylation analyses. S.Y. conducted gene expression analysis. R.J.E. performed the final polishing, final assembly cleanup, and KAT analysis. J.K. performed the genome annotation. R.J.E., E.K.F.C., and B.D.R. performed the *AMY2B* analyses. M.A.F. performed structural variance analyses. S.G. performed selection analysis. J.W.O.B. conducted the experimental analyses, M.E., T.T. and J.A.R. performed microbiome analysis. S.B., B.J. C. and B.M. performed biochemical analysis. M. B. performed metabolite analysis. S.Y. conducted statistical tests on biochemical and metabolome dataset. R.G.M, A.E.M., T.P.L.S. and M.T.P.G. commented on the manuscript. S.Y. and J.W.O.B. wrote the manuscript. All authors edited and approved the final manuscript.

