## Supplementary material for "Desert Dingo (*Canis lupus dingo*) genome provides insights into their role in the Australian ecosystem"

### 1 GENOME ASSEMBLY, ANNOTATION AND COMPARATIVE ANALYSES

#### 1.1 Sampling

The Desert Dingo Sandy dog was found in a remote region of South Australia in 2014. She was rescued with her two siblings and transported to eastern Australia. Subsequent genetic testing[1,2] showed she was a pure Desert Dingo.

#### 1.2 Pacific Bioscience Single Molecule Real-Time (SMRT) sequencing

Genomic DNA was prepared from skin biopsy. This was performed with supplemental RNase (Astral Scientific, Taren Point, Australia) and proteinase K (NEB, Ipswich, MA, USA) treatment, as per the manufacturer's instructions. Isolated gDNA was further purified using AMPure XP beads (Beckman Coulter, Brea, CA, USA) to eliminate sequencing inhibitors. DNA purity was calculated using a Nanodrop spectrophotometer (Thermo Fisher Scientific), and molecular integrity was assessed using pulse-field gel electrophoresis. DNA integrity was assessed by the Sage Science Pippin Pulse. A 0.75% KBB gel was run on the 9hr 10-48kb (80 V) program. DNA ladder used was the Invitrogen 1kb Extension DNA ladder (cat 10511-012). 150ng of DNA was loaded on the gel.

We generated two libraries that were size selected on Sage BluePippin gels (Sage Science, Beverly, MA, USA). Libraries were sequenced on Sequel machines with 2.0 chemistry recording 10 h movies. Sequencing was conducted at Arizona Genomic Institute, University of Arizona.

#### 1.3 Oxford Nanopore Technologies (ONT) PromethION sequencing

DNA (1 µg) was prepared for ONT sequencing using the 1D genomic DNA by ligation kit (SQK-LSK109, ONT) according to the standard protocol. Long fragment buffer was used for the final elution to exclude fragments shorter than 1000 bp. In total, 119 ng of adapted DNA was loaded onto a FLO-PRO002 PromethION flow cell and run on an ONT PromethION sequencing device (PromethION, RRID:SCR_017987) using MinKNOW (18.08.2) with MinKNOW core (v1. 14.2).

Base-calling was performed after sequencing with the GPU-enabled guppy basecaller (v3.0.3) using the PromethION high accuracy flip-flop model with config 'dna_r9.4.1_450bps_hac.cfg'. Sequencing was conducted at Kinghorn Centre for Clinical Genomics at the Garvan Institute of Medical Research, Sydney, Australia.

#### 1.4 10X Genomics Chromium sequencing

DNA was prepared following the protocol described above for SMRT sequencing. A 10X GEM library was barcoded from high-molecular-weight DNA according to the manufacturers recommended protocols. The protocol used was the Chromium Genome Reagent Kits v2 User Guide, manual part number CG00043 Rev B [3]. QC was performed using LabChip GX (PerkinElmer, MA, USA) and Qubit 2.0 Flurometer (Life Technologies, CA, USA). The library was run on a single lane of a v2 patterned flowcell. Paired-end sequencing with 150 bp read length was performed using the Illumina HiSeq X (Illumina HiSeq X Ten, RRID:SCR_016385) within the Kinghorn Centre for Clinical Genomics at the Garvan Institute of Medical Research, Sydney, Australia.

#### 1.5 Long read genome assembly

The SMRT and ONT reads were corrected and assembled with the Canu assembler (Canu, RRID:SCR_015880; v1.8.0) [4]. The assembled genome, with a total length of 2427850753 bp, consisted of 1834 contigs with an N50 length of 24.1 Mb (including 152 repeats of total length 16,671,837 bp) with no bubbles. There were 2,000,973 unassembled sequences of total length 13,107,822,345 bp. The resulting contigs were polished by aligning the raw reads to the assembly and correcting the sequencing errors using Arrow polishing [6]. There were 2,934,153 fixes implemented. Following Arrow polishing there were 1834 sequences with a total length of 2431109461 bp.

#### 1.6 Chromosome-length assembly using Hi-C data

Briefly, an *in situ* Hi-C library was prepared as previously described[5] from a blood sample from the same individual (www.dnazoo.org/methods). The Hi-C data was aligned to the polished contig set using Juicer[6], and used as input into the 3D-DNA pipeline[7] to produce a candidate chromosome-length genome assembly. We performed additional finishing on the resulting scaffolds using Juicebox Assembly Tools[8]. Fig 1b shows the contact matrices generated by aligning the Hi-C data set to the genome assembly before the Hi-C upgrade (on the left), and after Hi-C scaffolding (on the right). The matrices are visualised in Juicebox.js, a cloud-based visualisation system for Hi-C data [9] and are available for browsing at multiple resolutions at DNA Zoo [10]. This reduced the number of scaffolds to 210 (N50 64.2Mb), introducing 197 gaps. Subsequent polishing with Arrow closed 29 of these gaps, increasing contig N50 to 26.2 Mb.

#### 1.7 Gap filling, Pilon Polishing and Final cleanup

After scaffolding and correction, all raw SMRT and ONT reads were aligned to the assembly with Minimap2 (v2.16) (-ax map-pb/map-ont) and used by PBJelly (pbsuite v.15.8.24) [11] to fill gaps. It was able to completely close 74 gaps, increasing contig N50 to 36.2 Mb. A third round of Arrow polishing closed a further 25 gaps, increasing contig N50 to 40.7 Mb.

To further improve the assembly, another round of polishing was performed by aligning the 10X Chromium reads to the assembly using the linked-read analysis software provided by 10X Genomics, Long Ranger, v2.2.2[12]. Small indels were then corrected using Pilon v1.23 (diploid mode)[13].

The Pilon-polished genome was mapped onto CanFam v.31 chromosomes with PAFScaff v0.3.0 and underwent a final scaffold clean-up with Diploidocus v0.9.6 ("Nala" purge mode) to generate a high-quality core assembly, remove low-coverage artefacts and haplotig sequences, and annotate remaining scaffolds with potential issues. PacBio subreads (15.8 M subreads; 149.9 Gb) and ONT "pass" reads (6.12 M reads; 49.8 Gb) were mapped onto the assembly using Minimap2 v2.17 (-ax map-pb or -ax map-ont --secondary=no) [14] and read depth summaries calculated with BBMap v38.51 pileup.sh [15]. Any scaffolds with median coverage less than three (e.g., less than 50% of the scaffold covered by at least three reads) were filtered out as low-coverage scaffolds. Single-copy read depth was estimated using the modal read depth of 75X across the 5,736 single copy complete genes identified by BUSCO v3.0.2b [16]. This was used to set low-, mid- and high-depth thresholds for PurgeHaplotigs v20190612 [17] (implementing Perl v5.28.0, BEDTools v2.27.1 [18], R v3.5.3, and SAMTools v1.9 [19]) at 18X, 56X and 150X, to remove allelic contigs. PurgeHaplotig coverage parameter was adjusted to exclude gap regions and any scaffolds with ≥80% bases in the low/haploid coverage bins and ≥95% of their length mapped by PurgeHaplotigs onto another scaffold were filtered as haplotigs or assembly artefacts. Any other scaffolds with ≥80% low coverage bases were filtered as Low Coverage. In total, 11 sequences (93.2 kb) were removed as low-coverage artefacts and a further 31 (438.7 kb) were removed as probably haplotigs. Evaluation of the completion of the conserved single-copy genes was performed by by BUSCO v3.0.2b, short mode, implementing BLAST+ v2.2.31 [20], HMMer v3.2.1 [21], AUGUSTUS v3.3.2 [22] and EMBOSS v6.6.0) against Laurasiatheria_ob9 data set (n=6,253)

Following a second round of read mapping and depth filtering, no further scaffolds were identified for removal. The remaining 159 of the 201 Pilon-polished scaffolds were further classified based on read depth profiles and 51 scaffolds with <20% diploid coverage and ≥50% high coverage were marked as probable collapsed repeats. A single scaffold with "Diploid" depth as the dominant PurgeHaplotigs coverage bin and >50% match to another scaffold was marked as a possible repeat sequence[23].

Additional k-mer analysis of the final assembly was performed using KAT v2.4.2 [24] (Supplementary Table 1.2). KAT comp was used to compare k-mer frequencies from the 10x reads (16 bp barcode trimmed from read 1) with their copy number in the assembly. This comparison revealed no sign of missing data nor large duplications, including retention of haplotigs (Extended Data Fig. 2).

The genome was annotated using the homology-based gene prediction program GeMoMa (version 1.6.2beta)[25] and nine reference organisms including *Canis lupus familiaris*, *Vulpes vulpes*, *Felis catus*, *Sus scrof*, *Bos taurus*, *Ailuropoda melanoleuca*, *Ursus maritimus*, *Mus musculus*, and *Homo sapiens*. The assembled contigs were then aligned to CanFam3.1 for chromosome assignments[23].

Finally, Diploidocus v0.9.6 "vecpurge" mode (implementing blast+/2.9.0 tblastn) was used to screen the assembly for contaminants from the NCBI UniVec database (downloaded 05/08/2019) and the PacBio control sequence (MG551957.1). No additional scaffolds were masked, trimmed or purged. The estimated base error rate of the assembly is 0.00014 (Supplementary Information Table 1.2) and similar to other long-read genome [26].

#### 1.8 Mitochondrial genome assembly

The mitochondrion for Sandy was filtered out of the assembly at the initial haplotig purging step due to a high read depth. This 68.8 kb contig (tig00007654) was used as the basis for the mitochondrial chromosome. GABLAM v2.30.5 (implementing BLAST+ v2.9.0 blastn) mapped the 16,727 bp CanFam 3.1 mitochondrion onto tig00007654. The sequence was circularised by extracting the best complete match (positions 15268-31991) as the basis for the mitochondrial genome. Final Pilon polishing was performed by adding the mtDNA to the main Sandy assembly and mapping 10X Genomics linked reads using Long Ranger v2.2.2 before running Pilon v1.23 with the same settings as the main assembly. The 16,726 bp polished mitochondrial genome was then extracted and added back to the main nuclear genome assembly.

#### 1.9 Structural variation

To identify large structural differences in the dingo genome, structural variants from both Oxford Nanopore and PacBio sequence data were called relative to CanFam 3.1 and CanFam_GSD using a combination of minimap2 v2.17-r943-dirty[14], SAMTools v1.9[19], and sniffles v1.0.11[27].

#### 1.10 Genetic differentiation between the dingo and two domestic dog breeds

Mapping and quality control

To quantify the genetic differentiation between the dingo and the domestic dog breeds, we computed the pairwise Fst between the dingo and two domestic dog breeds, viz., the German shepherd dog (GSD) and the boxer. To perform this analysis, we obtained the previously published genomes from 10 dingoes[28], 20 GSDs and 14 boxers. These short read from these whole genome sequencing experiments were aligned against the de novo reference genome assembly of the dingo using the PALEOMIX pipeline[29]. As part of the PALEOMIX pipeline, adapter sequences were removed from the 3' ends of short reads using AdapterRemoval[30] v2.2.0, and low quality bases at the ends of reads (base quality < 2) were trimmed. Subsequently, bwa v0.7.16a was used to map the trimmed short reads to the de novo assembly of the dingo reference genome, using the backtrack algorithm of the PALEOMIX pipeline.

Genotype calling

Using the aligned reads from PALEOMIX, genotypes for the 44 samples were estimated using the HaplotypeCaller walker in Genome Analysis Toolkit (GATK v4.1.8.1). HaplotypeCaller was run using default parameters, but discarding bases with base quality lower than 15. Further, only variant sites were emitted during genotype calling, discarding all sites without any variation.

Computing Fst

Fst between each pair of populations, viz., dingo-GSD, dingo-boxer and GSD-boxer was computed using vcftools v0.1.16. The weighted version of the Weir Fst was computed across the genome in sliding windows of 100kb, with a step of 50kb. Variants with a minor allele frequency less than 0.05 in the combined dataset were discarded, as were sites with missingness greater than 20%. For the zoomed in version of the analyses focusing on *AMY2B* and *MGAM*, the exact same filters were used, but the sliding window size was reduced to 500 bp with a step of 100 bp.

#### 1.11 DNA methylome

DNA methylation data of GSD blood was downloaded from GSE136348. Dingo's blood DNA methylation library was sequenced on the Illumina HiSeq X platform (150 bp, PE), generating 281 million read pairs and yielding 14.5x sequencing coverage. Sequenced reads were trimmed using Trimmomatic[31] and mapped to the ASM325472v1 genome reference using WALT[32] with the following parameters: -m 10 -t 24 -N 10000000 -L 2000. The mappability of the MethylC-seq library was 85.36%. Duplicate reads were removed using Picard Tools v2.3.0. Genotype and methylation bias correction were performed using MethylDackel with additional parameters: --minOppositeDepth 5 --maxVariantFrac 0.5 --OT 20,148,20,120 --OB 25,145,25,145. The numbers of methylated and unmethylated calls at each CpG site were determined using MethylDackel (https://github.com/dpryan79/MethylDackel). Bisulphite conversion efficiency was 99.7%, estimated using unmethylated lambda phage spike-in control.

Segmentation of dingo's and GSD's blood DNA methylomes into CpG-rich unmethylated regions (UMRs) was performed using MethylSeekR[33] (segmentUMRsLMRs(m=meth, meth.cutoff=0.5, nCpG.cutoff=5, PMDs = NA, nCpG.smoothing = 3, minCover = 5).

To compare DNA methylation levels between proximal gene regulatory regions, we lifted over dingo TSS-associated UMRs to the GSD genome and GSD UMRs to the dingo genome. Next, we calculated average CpG methylation at UMRs and their corresponding lifted-over regions. UMRs showing more that 30% CpG methylation difference between dingo and GSD, were selected for the subsequent analysis. The TSS-associated UMRs correspond to transcriptionally permissive gene promoters in each genome.

To validate the difference in expression in *GAL3ST1* and *MAB21L1* we performed RT-qPCR on six dingoes and six GSDs. RNA was extracted from blood using TRI reagent protocol. Extracted total RNA was treated with DNase I Amplification Grade (Sigma). cDNA was prepared from RNA template in 20ul reaction mixture using ProtoScript cDNA synthesis kit (New England Biolabs, MA, USA). To analyse the results from RT-qPCR run, the comparative cycle threshold (Ct) method was used. The expression of *GAL3ST1* was quantified using following primer: GAL_F2 forward 5'-CTTGGCCCCGTTGTCCTCG-3' and GAL_F2 reverse 5'- TGACCGCAGAGGCAGCCT-3'. The expression of *MAB21L1* was quantified using following primers: MAB_F1 forward 5'-AGTGCATCTGGGCTCTTAGAC-3' and MAB_R1 reverse 5'- AACAAAAGTTGCGCTGAGACC-3'.

The RT-qPCR programme included an initial step of 95°C for 10min, followed by 40 cycles of 95° C for 10s and 60° C for 45s. To confirm that a single specific product was produced, amplification was followed by a melting curve from 60-95°C, rising by steps of 1°C. The gene expression was normalised using two housekeeping genes[34] *HNRNPH* (forward 5'-CTCACTATGATCCACCACG-3' and reverse 5'-TAGCCTCCATAACCTCCAC-3') and *GAPDH* (forward 5'-TGTCCCCACCCCCAATGTATC-3' and reverse 5'-CTCCGATGCCTGCTTCACTACCTT-3'). The unpaired t-test was performed to detect significance.

To assess the level of variation in the dingo genome, several approaches were employed to detect both small and large-scale variation. Small-scale variation generally smaller than 50 bases was detected in both the dingo assembly and CanFam_GSD relative to CanFam v3.1 using pairwise MUMmer4[35] (v4.0.0 beta 2) alignment databases.

#### 1.12 Tables

**Supplementary Information Table 1.1 Genome assembly and annotation statistics for the desert dingo assembly v CanFam3.1 and CanFam_GSD.**

| Statistic | Desert Dingo | CanFam3.1 | CanFam_GSD |
|---|---|---|---|
| Total sequence length | 2,349,862,946 | 2,410,976,875 | 2,407,291,559 |
| Total ungapped length | 2,349,829,267 | 2,392,715,236 | 2,401,147,102 |
| Number of contigs | 228 | 27,106 | 735 |
| Contig N50 | 40,716,615 | 267,478 | 20,914,347 |
| Contig L50 | 20 | 2,436 | 37 |
| Number of scaffolds | 159 | 3,268 | 429 |
| Scaffold N50 | 64,250,934 | 63,241,923 | 64,346,267 |
| Scaffold L50 | 14 | 15 | 15 |
| Number of gaps | 69 | 23,876 | 306 |
| BUSCO complete (genome) | 93.0% (91.7% single copy, 1.2% duplicate copy) | 92.2% (91.1% single copy, 1.1% duplicate copy) | 93.0% (91.6% single copy, 1.4% duplicate copy) |
| BUSCO fragmented (genome) | 3.6% | 4.0% | 3.67% |
| BUSCO missing (genome) | 3.4% | 3.8% | 3.4% |
| BUSCO complete (annotation) | 98.7% (96.7% single copy, 2.0% duplicate copy) | 95.1% (94.1% single copy, 1.0% duplicate copy) | 98.9% (96.5% single copy, 2.4% duplicate copy) |
| BUSCO fragmented (annotation) | 1.0% | 1.9% | 1.0% |
| BUSCO missing (annotation) | 0.3% | 3.0% | 0.1% |

**Supplementary Information Table 1.2: Estimates of error rate.**

| Assembly | k-mers unique to assembly | k-mers found in both | QV | Error rate |
|---|---|---|---|---|
| sandy_all.contigs | 1661265 | 2427814073 | 34.856 | 0.0003269 |
| sandy.all.contigs.arrow | 9760718 | 2431072781 | 37.177 | 0.0001915 |
| sandy.all.contigs.arrow.purged.HiC | 7864412 | 2347271088 | 37.964 | 0.0001598 |
| sandy.all.contigs.arrow.purged.HiC.arrow | 7612694 | 2347439679 | 38.106 | 0.0001546 |
| sandy.all.contigs.arrow.purged.HiC.pbjelly | 8906628 | 2350176975 | 37.428 | 0.0001807 |
| sandy.all.contigs.arrow.purged.HiC.pbjelly.arrow | 8269631 | 2350366783 | 37.751 | 0.0001678 |
| Final_assembly_GCA_003254725.2_ASM325472v2 | 7120099 | 2349824707 | 38.401 | 0.0001445 |

NOTE: Error rate was calculated in Merqury[36]. In brief, we estimate the probability P that a base in the assembly is correct as: $P = (\text{Kshared}/\text{Ktotal})^{1/k}$ where the Ktotal is the total number of k-mers found in an assembly and Kshared are the number of shared k-mers between the assembly and the read set. If the read set is assumed to completely cover the genome, any k-mer found only in the assembly ($\text{Kasm} = \text{Ktotal} - \text{Kshared}$) likely reflects a base error in the assembly consensus. Hence, the error rate E can be defined as: $E = 1 - P = 1 - (1 - \text{Kasm}/\text{Ktotal})^{1/k}$. QV is calculated as $-10 \times \log_{10}$ (Error rate) i.e. log-scaled probability of error for the consensus base calls.

**Supplementary Information Table 1.3 List of 44 unique genes detected in the regions overlapped with 132 identified structural variants.**

| |
|---|
| AHCYL2 |
| ATP11C |
| BPTF |
| CACNG1 |
| CCNB3 |
| CEP112 |
| CRB1 |
| DENND1B |
| ENSCAFE00000016325 |
| ENSCAFE00000422344 |
| ENSCAFG00000003879 |
| ENSCAFG00000044295 |
| GNE |
| HELZ |
| HTR1F |
| IKZF1 |
| JMJD8 |
| MCF2 |
| MED13L |

| |
|---|
| MGAM |
| MGAM2 |
| MOSPD1 |
| PIGQ |
| PRKCA |
| PSMD12 |
| RANBP17 |
| RNF38 |
| RNPC3 |
| SATL1 |
| SMKR1 |
| STRIP2 |
| TAS2R3 |
| UBR5 |
| ZMAT1 |
| ZNF654 |
| ZNF75D |
| ZPBP |

### 2 BIOCHEMICAL, PHYSIOLOGICAL AND MICROBIOME DIFFERENCES BETWEEN DINGOES AND GSDS

#### 2.1 Sample collection

Biochemical studies were performed on 17 dingoes two different dingo sanctuaries and 15 GSDs from two different Kennels in December 2018. Of these 14 were from Bargo Dingo Sanctuary (7 males and 7 females) and 3 were from Pure Dingo Sanctuary (1 male and 2 females) in south east New South Wales (NSW), Australia. All dingoes were pure as determined by microsatellite testing [1,2]. The age was 3.8 ± 0.44 (SE) years. Five dingoes were born in the wild but humanized before 6 weeks of age. The remaining 11 dingoes were sanctuary born. All had daily interactions with humans. No consistent differences were observed between these groups that would suggest results were influenced by whether dingoes were wild versus sanctuary born did the influence results. One male dingo from Bargo was excluded from all analyses as his partner suffered a broken leg prior to the study and his feeding behaviour was modified. The dingoes were housed in mated pairs and were not kept as pets. Volunteers fed and walked the dingoes daily. The dingoes were socialized but rarely travelled from the sanctuary.

Fifteen GSDs included in the study were tested in December 2018. Eleven were from Kingvale Kennels (3 males and 8 females) and four were Allendell Kennels (1 male and 3 females). Both Kennels were in south east, NSW Australia. The mean age was 3.66 ± 0.44 years. Two female GSD's were subsequently excluded from the study as they came into oestrus within 10d of the study. All GSD's were registered with the Australian Kennel Club and showed no evidence of any genetic diseases including hip-dysplasia. The GSD's were kept in large runs with fewer males than females in each kennel. All GSD's were socialized and used to travelling distances in cars and trailers.

No obvious differences were observed between the maintenance conditions of the dingo sanctuaries and the domestic dog kennels and results were pooled for analysis.

#### 2.2 Experimental diets and treatments

To avoid a possible bias in the results due to diet differences between individuals of kennels and sanctuaries, we standardised the diet of dingoes and GSDs for 14d. Canids were fed throughout the evening on standard "Blackhawk" kibble and raw chicken for the first 10d. From day 11, canids were transitioned to rice and chicken food so they were fed rice and chicken only on day 14. Fresh untreated rainwater was transported to all Sanctuaries and kennels.

On the evening of day 1, all canids were treated for fleas, ticks and worms with Advocate for dogs (Bayer) and given the antibiotic Neo-Sulcin (Jurox Animal Health) in kg/ dependent doses. On days 2 and 3 they were then given the probiotic Protexin (Protexin Veterinary) to recolonise the gut microbiota in kg/ dependent doses. Blood was drawn in the morning of day 15 for biochemical and were stored at 4°C for metabolism studies.

#### 2.3 Amylase copy number

We used droplet digital PCR (ddPCR) to directly quantify DNA copy number variation [37]. ddPCR was performed using a QX100 ddPCR system (Bio-rad). Each reaction was performed in a 20µl reaction volume containing 10µl of 2x ddPCR Supermix (Bio-rad), 1µl of each 20x primer/probe, 1µl of DraI restriction enzyme (New England BioLabs #R0129S),

5μl of DNA template (4ng/μl) and 2μl $ddH_2O$. Copy number data were rounded to the nearest whole number and presented as copies per individual chromosome. Primer sequence for Amy2B: forward 5′-CCAAACCTGGACGGACATCT-3′ and reverse 5′-TATCGTTCGCATTCAAGAGCAA-3′ with FAM probe: 6FAM–TTTGAGTGGCGCTGGG-MGBNFQ. Primer sequence for C7orf28b-3: 5′-GGGAAACTCCACAAGCAATCA-3′ and reverse 5′-GAGCCCATGGAGGAAATCATC-3′ with HEX probe HEX-CACCTGCTAAACAGC-MGBNFQ. Statistical significance in amylase copy number difference and biochemical studies were analyzed by simple t-tests using Prism software program version 8.0 (www.graphpad.com).

#### 2.4 Serum metabolites

Amylase, cholesterol, triglycerides, and lipase
We tested for metabolic differences associated with starch digestion and fat metabolism [38]. Amylase, cholesterol, triglycerides, and lipase and were assayed using the Thermo Scientific Konelab Prime 30i at the Veterinary Pathology Diagnostic Services Laboratory (VPDS), University of Sydney. Statistical significance was determined as described above.

Serum lipoprotein profile analysis
Serum was fractionated on an AKTA FPLC system (GE Healthcare Life Sciences) using two Superdex 200 columns (GE Healthcare Life Sciences) connected in series. Plasma (200μL) was loaded onto the columns, which had been pre-equilibrated with PBS (10mM $NaH_2PO_4$, 137mM NaCl, 2.7mM KCl, pH 7.4). Lipoproteins were separated at a flow rate of 0.25 mL/min. Fractions were collected at 1min intervals and immediately analysed on an AU480 Auto-Analyser (Beckman Coulter) for total cholesterol levels using the Wako Cholesterol E reagent (Wako Diagnostics).

Bile acid quantification analysis
To examine whether differences in cholesterol levels influences bile acid production in dingoes and GSDs, we quantified free bile acids in canine plasma using a liquid chromatography tandem mass spectrometry (LCMS/MS) assay[39]. We measured concentration of two primary bile acids: Cholic acid (CA) and Chenodeoxycholic acid (CDCA) and three secondary bile acids: Ursodeoxycholic acid (UDCA), Deoxycholic acid (DCA) and Lithocholic acid (LCA). Standards for all bile acids were prepared at 0, 10, 20, 40, 60, 100, and 200nM concentration from a 1uM combined stock solution. Deuterium labelled standards; $d_4$CA, $d_4$DCA, $d_4$CDCA and $d_4$LCA were combined to a final concentration of 4μM and used as internal standards to correct for variability and losses during processing, a 10ul aliquot of IS was added to each calibrator (final volume 200μL). Each canine plasma sample (30-100μL available for testing) was mixed with 10μl aliquot of combined deuterated IS and 4 volumes of acetonitrile. The mixture was vortexed, centrifuged at 10,000rpm for 10min in order to remove proteins. The supernatant was transferred to a clean tube, and vacuum dried before reconstitution in a 50:50 solution of methanol and water (200μL). The sample was filtered into reduced volume vials and ready for LCMS/MS analysis.

The ultra-performance liquid chromatography MS detector (UPLC-MSD) assembly consisted of an Accela AS injector, Accela UPLC pump, and a TSQ Vantage bench-top mass spectrometer (ThermoFisher Scientific, Waltham, MA) fitted with a heated electrospray probe (HESI). Solutions of the tested five bile acids; including labelled analogs (200μM in 50% methanol) were infused at 10μl/min by means of a syringe pump into the detector. Collision induced dissociation (CID) experiments in negative ion mode were carried out to determine the parameters at which optimum sensitivity was achieved for these metabolites.

We found that bile acids under CID; either formed no useful fragments for quantitation or rendered deprotonated molecular ions. The found selected-reaction monitoring (SRM) transitions were then set in the method mass spectrometry detector parameters. Mass spectra were accumulated during 0.2s per SRM. Capillary voltage, capillary temperature and collision gas pressure (Argon) were set to 3000V, 300°C and 1.0Torr. respectively. Sheath and auxiliary gas valves (nitrogen) were set at 20 and 10 arbitrary units.

Standards and samples (20µL) were injected into a Waters Acquity BEH18 column (100mm x 2.1mm x 1.7µm) heated at 40°C. The binary solvent gradient consisted of 5mM ammonium formate (mobile phase A) and acetonitrile (mobile phase B) at a constant flow rate of 200µL/min. Initial solvent composition at injection was 40% B, followed by a 5min gradient to 50% B and a fast gradient ramp to 80% B (1min), B was increased again to 95% (2min), held for 4min and then reverted to initial conditions (0.1min) for equilibration, with a total run time of 18min. The column flow was directed into the MSD. Retention times and mass transitions are shown in Supplementary Information Table 2.2; differences in retention times were observed between plasma samples likely due to sample matrix components.

Calibration curves for each individual bile acid were plotted using the peak area ratios (Y axis; peak area of the bile acid divided by peak area of its corresponding deuterated counterpart, UDCA was normalised to $d_4$-CDCA) versus nM standards concentration (X axis). All spectra were processed and peak areas integrated using Xcalibur™ software (version 2.2, 2011, ThermoFisher Scientific, Waltham, MA). Automated data processing was performed using the LCQuan feature of the software. The concentrations of the endogenous metabolites in the sample extracts were obtained from these calibration curves and calculated using dilution factors. To test significant differences in the individual bile acid concentration t-tests were performed on individual bile acids using Prism software program version 8.0 (www.graphpad.com) with two outliers removed for dingo and two for GSD using ROUT method of Prism and a false discovery rate (Q) of 1%.

#### 2.5 Microbiome analysis

Simultaneously with the biochemical studies, scat from same set of dingoes and GSD's were sampled on day 1 and 14. Stool samples for microbiome analysis were collected in the morning, taking care not to pick up contaminants such as soil and plant material. The samples were placed into sterile bags and kept cold until they reached the laboratory, where they were immediately stored at -80$^0$C until assayed.

DNA was extracted from thawed stool samples (0.3g) using the Qiagen Powersoil kit according to the manufacturer's instruction (cat# 1288-100; Hilden, Germany). However, instead of vortexing, samples were subjected to physical lysis in a bead-beater (TissuerLyzer11, Qiagen) for three min at 30Hz. DNA was eluted in molecular grade water and stored at -80$^0$C. The V3-V4 region of the 16S rRNA gene were amplified and sequenced as described in[40]. Library preparation and pair end sequencing was performed (2x300 cycles) on the Illumina MiSeq platform at the Ramaciotti Centre for Genomics (UNSW, Sydney, Australia).

16S rRNA gene sequence data were quality filtered and trimmed using TRIMMOMATIC version 0.36 truncating reads if the quality was below 12 in a sliding window of 4bp. USEARCH version 10.0.240 [41] was used to merge and quality filter sequencing reads between 350 and 500 nucleotides. Sequences that appeared less than 8 times were removed. Processed reads were then concatenated into a single file and subsequently dereplicated to form unique sequences. Unique sequences were clustered into zero-radius operational taxonomic units (zOTUs) using the UNOISE3 algorithm implemented in USEARCH [41]. Chimeras were removed in reference mode using UCHIME together with the SILVA SSURef NR99 database version 132 [42].

The zOTU sequences were taxonomically classified using BLASTn alignments against the SILVA database. zOTUs without any taxonomic assignment were removed from the dataset. No zOTUs were found to be assigned to chloroplast in the dataset. The number of final zOTUs was 9951. Data were visualized using the ggpubr package[43]. For alpha diversity measures, each sample was subsampled 100 times to a count of 164,700 counts per sample and the average was taken. zOTU richness and diversity indices (Shannon, ACE and Chao1) were calculated in R (version 3.6.0) using the vegan package[44] and statistically compared between Dingoes and GSD using the Wilcoxon Rank Sum test.

For beta diversity, the rarefied data were square root transformed. Bray-Curtis distances were calculated and visualized on a non-metric multi-dimensional scaling (NMDS) plot. The zOTU sequences were aligned using MAFFT [45] and a phylogenetic tree was calculated using FastTree [46] to calculate weighted UniFrac distances, which were visualized on a principal coordinate analysis (PCoA) plot. Differences between in the beta diversity of dingoes and GSD communities were analyzed using a pairwise.adonis test (https://github.com/bwemheu/pairwise.adonis).

Metabolic potential of the microbial community was evaluated using the predictive metagenomic analysis tool, Tax4Fun2[47]. Relative abundances of taxa or predicted functions were examined using the phyloseq package[48]. Significant differences of microbial taxa between the canids were analyzed at phylum, family and genus levels using the Analysis of the Composition of the Microbiome (ANCOM; v2.0) [49]. ANCOM evaluates statistical significance of the taxa or predicted functions using log-ratio transformed data. Benjamini-Hochberg correction was applied to correct for multiple comparison testing and a p value of less than 0.05 was considered statistically significant.
Bacterial composition of the dingo and German Shephard dog were visualized with bar plots for each location of sampling, using phyloseq R package version 1.32.0[48]. We visualized the relative abundance of top ten most abundant OTUs. We analyzed the differential abundance of the OTUs, using metacoder R package version 0.3.4[50]. Prior to differential abundance analysis were taxonomy concatenated according to Species, using phyloseq. The $\log_2$ ratio of median proportions of relative abundance between the dingo and German Shephard dog and significance were determined using a Wilcoxon rank-sum test followed by a Benjamini–Hochberg (FDR) correction for multiple comparisons. The $\log_2$ ratio of median proportions of insignificant different abundant taxon were removed for clarity.

#### 2.6 Scat metabolite analysis

Following Carthey et al. [51] with modifications, we chose solid phase microextraction (SPME) gas chromatography-mass spectrometry (GC-MS) to sample and analyse the volatile organic compounds (VOC's) in the headspace above scats. Sample vials were incubated for 5min at 60°C, before a pre-conditioned 50/30 µm divinylbenzene / carboxen / polydimethylsiloxane SPME fibre (Catalog # 57298-U, Supelco, Bellafonte, PA) was exposed to the headspace for 25min (60°C).

The fibre was desorbed in the inlet port for 2min, then purged with carrier gas in the inlet at a split flow of 50mL/min for 23 min, to condition it for the next analysis. The GC was fitted with a high-polarity free fatty acids phase column (HP-FFAP, 50mm × 0.2mm, 0.3µm film thickness, Agilent Technologies) and helium was used as the carrier gas (1mL/min). The oven temperature program was 40°C for 2 min, then 5°C per min increase to 240°C where it was held for 10min. The mass spectrometer was operated in electron impact ionization (EI+)

mode, the ion source temperature was 200°C and the GC transfer line temperature was 245°C.

Parameters for the AMDIS analysis are listed in Table S4.3. This deconvolution step was included in order to obtain pure mass spectra for VOC's where chromatographic peaks were overlapping (near co-eluting) with those from other compounds. The deconvolution also served to remove background/baseline signals from the data. Deconvoluted mass spectra were automatically searched against the Wiley 9/NIST 2011 Combined (de-duplicated) Library of Mass Spectra by the AMDIS algorithm. All library matches with a match factor equal to, or exceeding 85/100 were considered positive putative peak identifications. A new target compound library (*.msl) was manually curated from putative VOC identifications and chromatographic retention times (RT's) obtained from individual AMDIS analyses of five dingo and five GSD scat (single replicate/scat) data files. Mass spectra for these target VOC's were copied from the NIST/Wiley Library of mass spectra to the target library. Compounds that were also found during analysis of control (empty) vials were excluded from the target library. These were mainly siloxanes and plasticizers from the vials, septa, column and SPME fiber. The final target library contained 57 unique compounds, following manual removal of duplicate entries that showed different names for the same compound.

MS data for all pseudo replicates of all scats was re-analysed with AMDIS using its batch process function, referencing the new target compound library and searching within a non-penalized RT window of ±0.2 min of target compound library RT values. Results were saved to a tab-delimited text file. This file listed the target compounds found in each scat pseudo-replicate, along with various measures of compound signal intensity and other data metrics. The AMDIS output was used as an input for Metab R package[52] and were analysed by using time window difference of 0.2 min. Base peak was used as a preferred measure of signal intensity output from the AMDIS deconvolution. A t-test for each chemical was carried out in R to identify significant difference between dingo and domestic dog. To confirm the identity of compounds from the list of putative identities generated by AMDIS's NIST / Wiley library search authentic standard for VOC's of interest were analysed by smearing a small amount on the inside of headspace vials, using a cotton bud. The significant VOC's were confirmed by both retention time similarity and library matching of mass spectra.

#### 2.7 Tables

**Supplementary Information Table 2.1 Details of dingoes and GSD's included in the study**

| Canid | Code | Locality | Age | Sex |
|---|---|---|---|---|
| Dingo | W0378 | Bargo | 3 | F |
| Dingo | W0235 | Bargo | 4 | M |
| Dingo | X3171 | Bargo | 8 | F |
| Dingo | W0379 | Bargo | 3 | F |
| Dingo | W0303 | Bargo | 3 | M |
| Dingo | X3172 | Bargo | 8 | M |
| Dingo | W0381 | Bargo | 4 | F |
| Dingo | W0382 | Bargo | 1.5 | F |
| Dingo | X1039 | Bargo | 3 | M |
| Dingo | W0234 | Bargo | 4 | F |
| Dingo | W0380 | Bargo | 5 | M |
| Dingo | W0363 | Bargo | 1 | M |
| Dingo | W0383 | Bargo | 3 | F |
| Dingo | X3170 | Bargo | 8 | M |
| Dingo | W0296 | Pure Dingo | 4 | F |
| Dingo | W0329 | Pure Dingo | 4 | M |
| Dingo | W0330 | Pure Dingo | 4 | F |
| GSD | GSD02 | Allendell | 4 | F |
| GSD | GSD03 | Allendell | 2 | M |
| GSD | GSD04 | Allendell | 4 | F |
| GSD | GSD05 | Allendell | 6 | F |
| GSD | GSD06 | King Vale | 1. 8 | F |
| GSD | GSD16 | King Vale | 5.9 | M |
| GSD | GSD07 | King Vale | 1.8 | F |
| GSD | GSD08 | King Vale | 3.6 | F |
| GSD | GSD09 | King Vale | 5.6 | F |
| GSD | GSD10 | King Vale | 5.9 | F |
| GSD | GSD11 | King Vale | 2.2 | M |
| GSD | GSD12 | King Vale | 2 .1 | F |
| GSD | GSD13 | King Vale | 2.3 | F |
| GSD | GSD14 | King Vale | 5 .6 | M |
| GSD | GSD15 | King Vale | 4 .1 | F |

Relative to the University of New South Wales Bargo Dingo Sanctuary is 94.9 km SW and Pure Dingo is 102km North. Kingvale Kennels is 10km west of Bargo Dingo Sanctuary and Allendell Kennels 38km north.

**Supplementary Information Table 2.2 The *m/z* transitions for quantitation and retention times of five tested bile acids. (*) Retention time shift in plasma**

| Bile acid | Retention time Stds (sample) | Collision Energy (EV) | *m/z* Transition |
|---|---|---|---|
| Cholic acid (CA) | 5.76 (4.74)* | 40 | 407/391,345/289 |
| d4-Cholic acid | 5.75 (4.69)* | 40 | 411/411 |
| Ursodeoxycholic acid (UDCA) | 6.52 (5.43)* | 35 | 391/391 |
| Chenodeoxycholic acid (CDCA) | 9.34 (9.21)* | 35 | 391/391 |
| d4-Chenodeoxycholic acid | 9.35 (9.21)* | 35 | 395/395 |
| Deoxycholic acid (DCA) | 9.53 (9.42)* | 35 | 391/345 |
| d4-Deoxycholic acid | 9.52 (9.39)* | 35 | 395/395 |
| Lithocholic acid (LCA) | 11.63 | 5 | 375/375 |
| Bile acid | 11.66 | 5 | 379/379 |

**Supplementary Information Table 2.3 Total number of quality filtered reads for each group for microbiome analysis**

| Group | Total reads for group | Median | Range |
|---|---|---|---|
| Dingo | 4,258,047 | 202,391 | 183,872 – 238,343 |
| GSD | 3,573,342 | 205,911 | 178,529 – 213,854 |

**Supplementary Information Table 2.4 Analysis of the Composition of the Microbiome (ANCOM). Analysis results for phylum, family and genus level at relative abundance of taxa. The level of significance cutoff = 0.05 with Benjamini-Hochberg correction.**

| **Taxa id** | **Dingo (%)** | **GSD (%)** | **Enriched** |
|---|---|---|---|
| **Phylum** | | | |
| Tenericutes | 0.006 | 0.013 | GSD |
| **Family** | | | |
| Acidaminococcaceae | 0.142 | 1.887 | GSD |
| Anaeroplasmataceae | 0.006 | 0.013 | GSD |
| Burkholderiaceae | 0.080 | 0.848 | GSD |
| Clostridiaceae_1 | 10.342 | 1.805 | Dingo |
| Corynebacteriaceae | 0.000 | 0.088 | GSD |
| Family_XIII | 0.000 | 0.617 | GSD |
| Lactobacillaceae | 0.777 | 1.166 | GSD |
| Leuconostocaceae | 0.009 | 0.606 | GSD |
| Marinifilaceae | 0.000 | 0.036 | GSD |
| Muribaculaceae | 0.100 | 0.691 | GSD |
| Peptococcaceae | 0.136 | 0.681 | GSD |
| Prevotellaceae | 9.863 | 19.237 | GSD |
| Rikenellaceae | 0.003 | 0.170 | GSD |
| Ruminococcaceae | 0.657 | 1.804 | GSD |

| | | | |
|---|---|---|---|
| Streptococcaceae | 0.178 | 2.410 | GSD |
| Succinivibrionaceae | 0.071 | 1.082 | GSD |
| Tannerellaceae | 0.021 | 0.226 | GSD |
| **Genus** | | | |
| *[Eubacterium]_brachy_group* | 0.000 | 0.465 | GSD |
| *[Eubacterium]_hallii_group* | 0.008 | 0.020 | GSD |
| *[Eubacterium]_nodatum_group* | 0.000 | 0.147 | GSD |
| *[Ruminococcus]_gnavus_group* | 1.644 | 0.474 | Dingo |
| *[Ruminococcus]_torques_group* | 0.288 | 0.715 | GSD |
| *Allisonella* | 0.000 | 0.039 | GSD |
| *Alloprevotella* | 3.565 | 9.674 | GSD |
| *Anaerobiospirillum* | 0.070 | 0.524 | GSD |
| *Anaeroplasma* | 0.006 | 0.013 | GSD |
| *Butyricicoccus* | 0.033 | 0.093 | GSD |
| *Catenisphaera* | 0.008 | 0.647 | GSD |
| *Citrobacter* | 0.041 | 0.397 | GSD |
| *Clostridium sensu stricto 1* | 10.342 | 1.805 | Dingo |
| *Corynebacterium* | 0.000 | 0.088 | GSD |
| *Enterobacter* | 0.027 | 1.273 | GSD |
| *Enterorhabdus* | 0.009 | 0.027 | GSD |
| *Faecalibacterium* | 0.179 | 0.661 | GSD |
| *Faecalitalea* | 0.052 | 0.022 | Dingo |
| *Flavisolibacter* | 0.005 | 0.000 | Dingo |
| *Fournierella* | 0.010 | 0.137 | GSD |
| *GCA-900066575* | 0.000 | 0.007 | GSD |
| *Hafnia-Obesumbacterium* | 0.000 | 0.024 | GSD |
| *Howardella* | 0.005 | 0.024 | GSD |
| *Klebsiella* | 0.001 | 0.071 | GSD |
| *Kluyvera* | 0.001 | 0.113 | GSD |
| *Lachnospira* | 0.016 | 0.224 | GSD |
| *Lactobacillus* | 0.776 | 0.836 | GSD |
| *Lactococcus* | 0.004 | 2.132 | GSD |
| *Leuconostoc* | 0.009 | 0.606 | GSD |
| *Negativibacillus* | 0.003 | 0.199 | GSD |
| *Oribacterium* | 0.007 | 0.141 | GSD |
| *Oscillospira* | 0.001 | 0.021 | GSD |
| *Paeniclostridium* | 0.613 | 0.075 | Dingo |
| *Pantoea* | 0.000 | 0.079 | GSD |
| *Parabacteroides* | 0.021 | 0.226 | GSD |
| *Parasutterella* | 0.009 | 0.036 | GSD |
| *Pediococcus* | 0.001 | 0.331 | GSD |
| *Peptococcus* | 0.136 | 0.681 | GSD |
| *Phascolarctobacterium* | 0.142 | 1.887 | GSD |
| *Plesiomonas* | 0.000 | 0.060 | GSD |
| *Prevotellaceae Ga6A1 group* | 0.239 | 2.611 | GSD |

| | | | |
|---|---|---|---|
| *Raoultella* | 0.000 | 0.008 | GSD |
| *Rikenellaceae RC9 gut group* | 0.003 | 0.170 | GSD |
| *Ruminiclostridium 9* | 0.000 | 0.011 | GSD |
| *Ruminococcaceae UCG-005* | 0.143 | 0.428 | GSD |
| *Sellimonas* | 0.056 | 0.168 | GSD |
| *Succinivibrio* | 0.000 | 0.558 | GSD |
| *Sutterella* | 0.032 | 0.716 | GSD |
| uncultured_f_Muribaculaceae | 0.100 | 0.691 | GSD |
| uncultured_f_Eggerthellaceae | 0.001 | 0.008 | GSD |
| uncultured_f_Peptostreptococcaceae | 3.326 | 0.556 | Dingo |

**Supplementary Information Table 2.5 GC-MS parameters used for AMDIS deconvolution of the data to identify target volatile organic compounds (VOC's) and to produce signal intensity measurements for the target VOC's in each scat pseudo-replicate**

| Parameter | Value | units | notes |
|---|---|---|---|
| Minimum match factor | 85 | /100 | (Identif. Tab↓) |
| Multiple identifications per compound | Not enabled | | |
| Show standards | Not enabled | | |
| Only reverse search | Not enabled | | |
| Type of analysis | Simple | | To create target library |
| | Use RT | | To analyse data referencing target comound library |
| RT Window | 0.2 | minutes | As above |
| Match Factor Penalties level | Average | | As above |
| Low m/z, high m/z | Auto | | (Instr. Tab↓) |
| Use Scan Sets | Not enabled | | |
| Threshold | Off | | |
| Scan Direction | None | | |
| Instrument Type | Quadrupole | | |
| Data File format | NetCDF | | |
| Component width | 20 | Scans | (Deconv. Tab↓) |
| Omit m/z | 73, 207, 281, 267, 341 | m/z | Column siloxanes |
| Adjacent Peak Subtraction | Two | Peaks | |
| Resolution | Medium | | |
| Sensitivity | High | | |
| Shape requirements | Low | | |
| Target Compounds Library | Blank | | Initial analysis (Libr. Tab) |
| Target Compounds Library | Scat VOC's | | For final analysis of all data |
| Column bleed | 207 | m/z | (QA/QC Tab) |
| Filters | Not enabled | | |

#### EXTENDED FIGURES

##### Extended Data Fig. 1: Desert dingo genome data generation summary

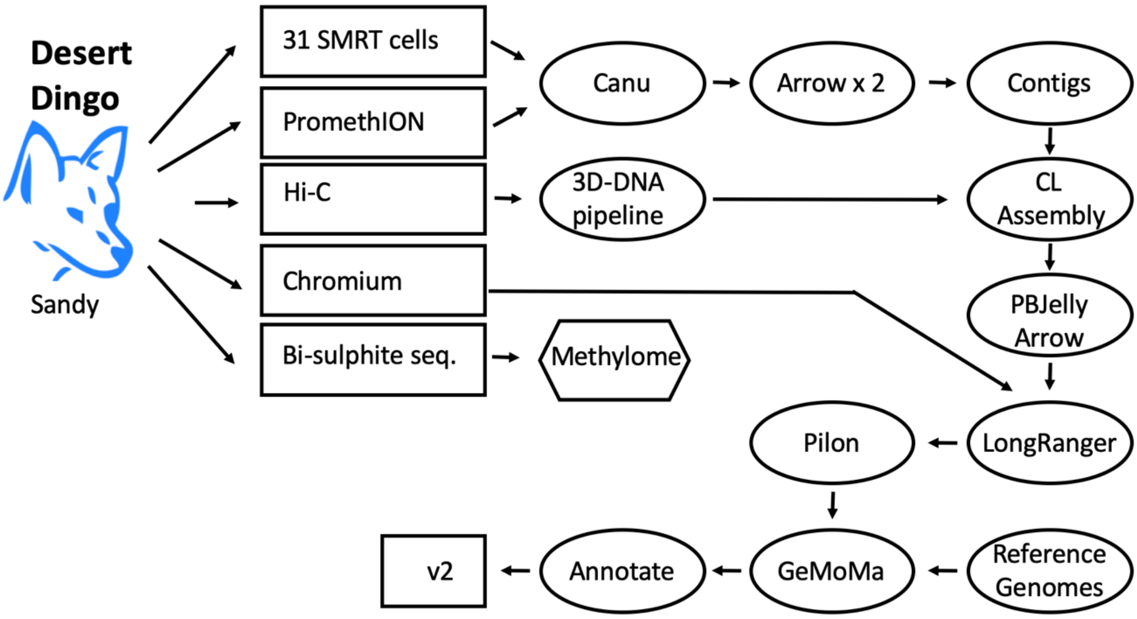


The desert dingo Sandy was found in a remote region of South Australia in 2014. Subsequent genetic testing showed she was a pure Desert Dingo. Initial PacBio sequences were generated from 34µg of high quality HMW DNA that was shipped to the Arizona Genome Institute, University of Arizona. The HMW DNA was bead purified and processed with 15kb blue pippin size selection to construct 2.3µg of high quality PacBio library. A total of 31 productive SMRT cells produced a total of 150.11Gb (4.84Gb/cell) with an average read length N50 of 17kb. Additional long read data was gathered from the Garvan Institute Oxford Nanopore PromethION instrument (guppy bascaller Version 3.0.6+9999d81) to ~85X genome coverage, each, based on a genome size estimate of 2.4Gb. All long read sequences were assembled with the Canu v1.8 algorithm then error corrected twice using the Arrow genomic consensus polishing module. This assembly was scaffolded to chromosome-length by the DNA Zoo following the methodology described here: www.dnazoo.org/methods. Briefly, an *in situ* Hi-C library was prepared from a blood sample from the same animal. The Hi-C data was processed using Juicer, and used as input into the 3D-DNA pipeline to produce a candidate chromosome-length (CL) genome assembly. We performed additional finishing on the scaffolds using Juicebox Assembly Tools. The assembly was then long-read gap filled with the PBJelly algorithm, and the additional data error corrected using Arrow. The Chromium 10X data obtained at the Garvan Institute the was mapped onto the assembly with the Long Ranger v2.1.6 program and the final assembly was then polished using the Pilon algorithm. The assembly has a size of 2.35 Gb, consists of 159 scaffolds with a contig and scaffold N50 length of 64.3 Mb (contig L50=20, scaffold L50=14) and 33.7 kb of gap sequence. The genome was annotated using the homology-based gene prediction program GeMoMa (version 1.6.2beta) and nine reference organisms. The assembled contigs were then aligned to CanFam3.1 for chromosome assignments.

**Extended Data Fig. 2: KAT kmer analysis of Sandy desert dingo assembly.**

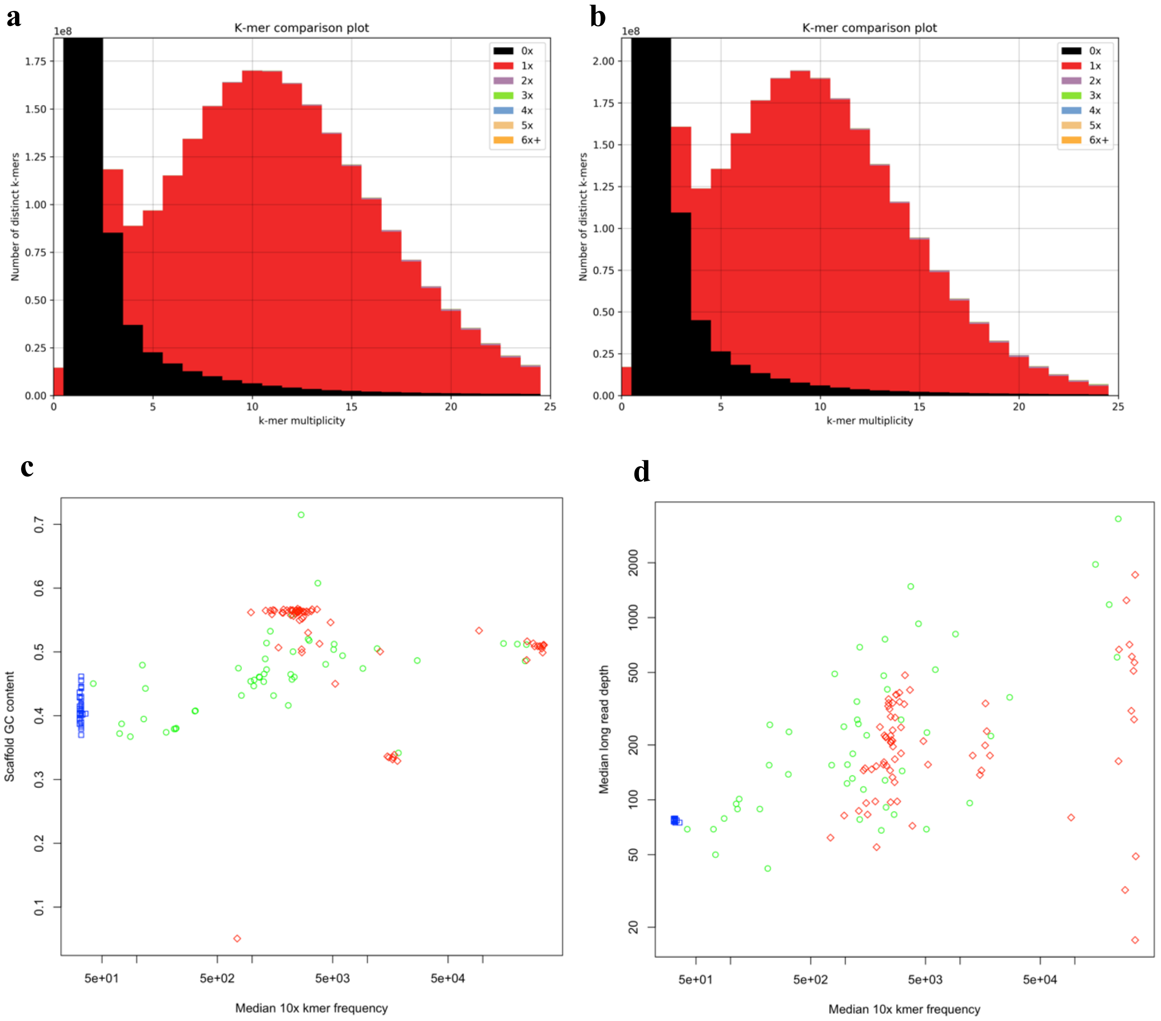


10x read kmers frequency distributions for kmers with different assembly copy numbers derived from **a,** Read 1 (16bp barcodes trimmed) and **b,** Read 2 (barcodes not trimmed). **c,** Scaffold GC content plotted against median 10x kmer frequency. Blue square: main chromosome; green circle: placed scaffold; red diamond: unplaced scaffold; cross: suspect scaffold. **d,** Median long read depth versus Median 10x kmer frequency.

**Extended Data Fig. 3: Synteny plot of Chromosome 16. Comparing CanFam3.1 vs CanFam_GSD.**

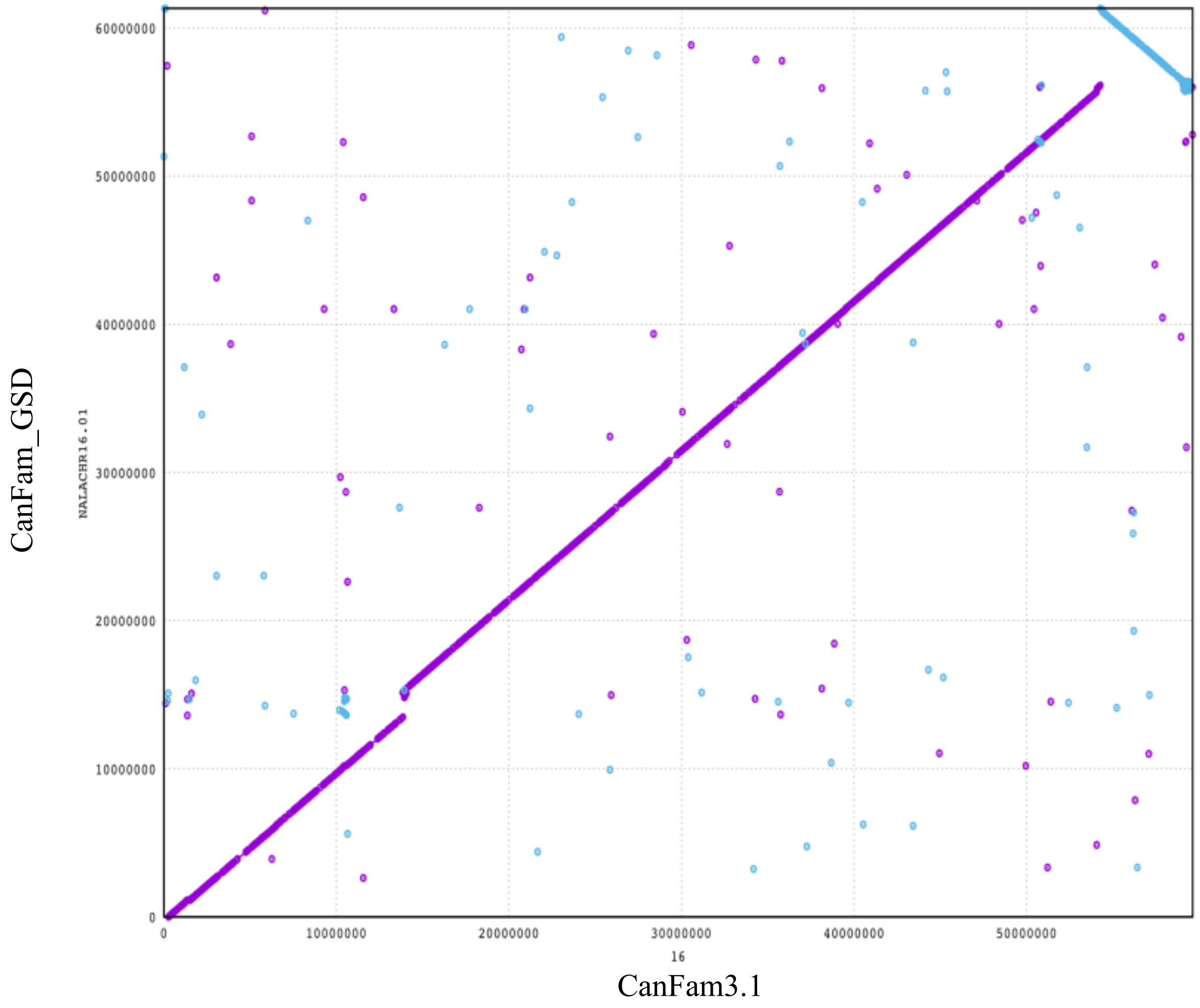


CanFam_GSD contains a single large inversion and a large insertion relative to CanFam3.1.

**Extended Data Fig. 4:** Selection scan focusing on *AMY2B*, *MGAM1* and *MGAM2* with the sliding window of 500 bp with a step of 100 bp.

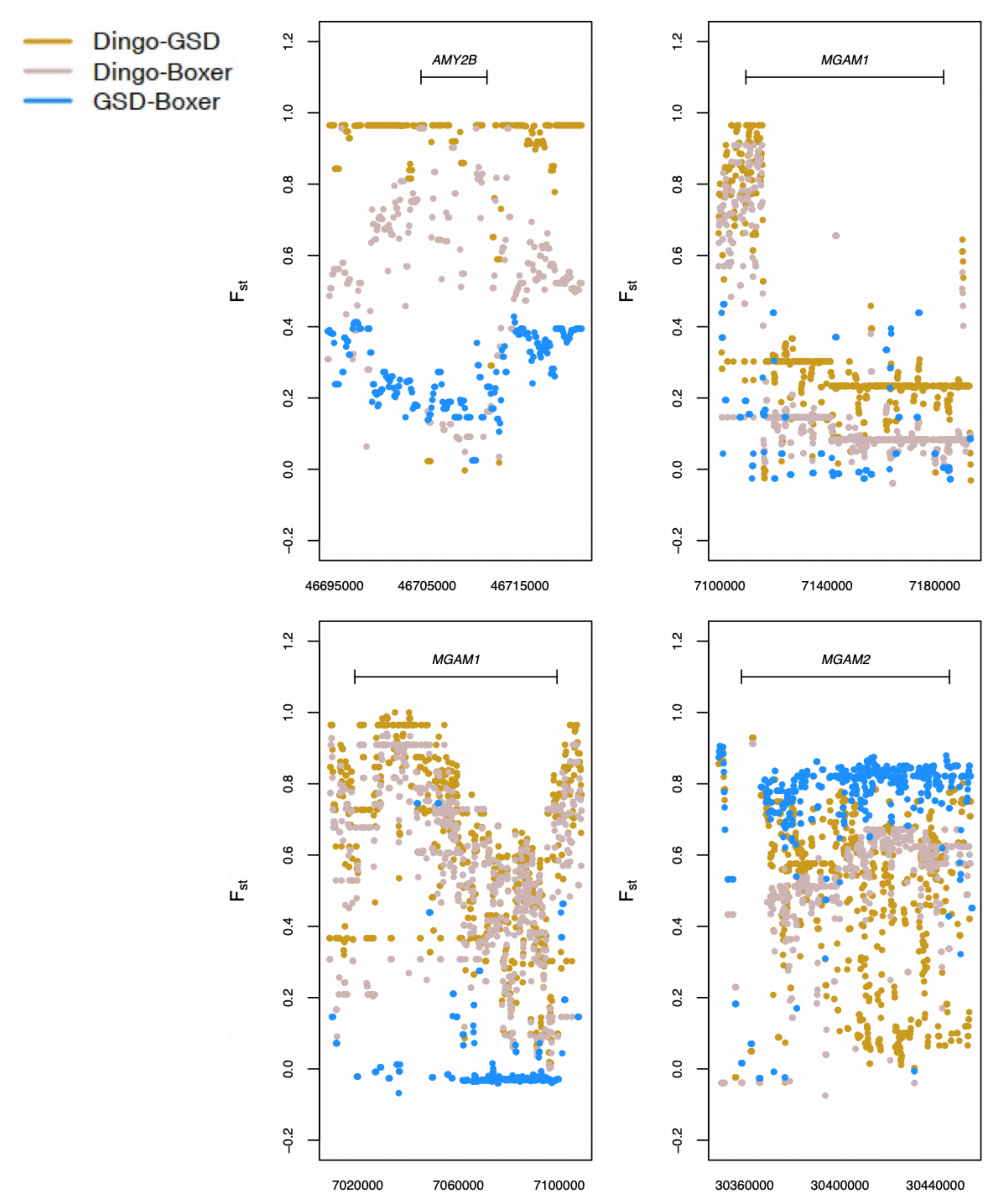

**Extended Data Fig. 5: DNA Methylome comparison between the dingo and German Shepherd dog.**

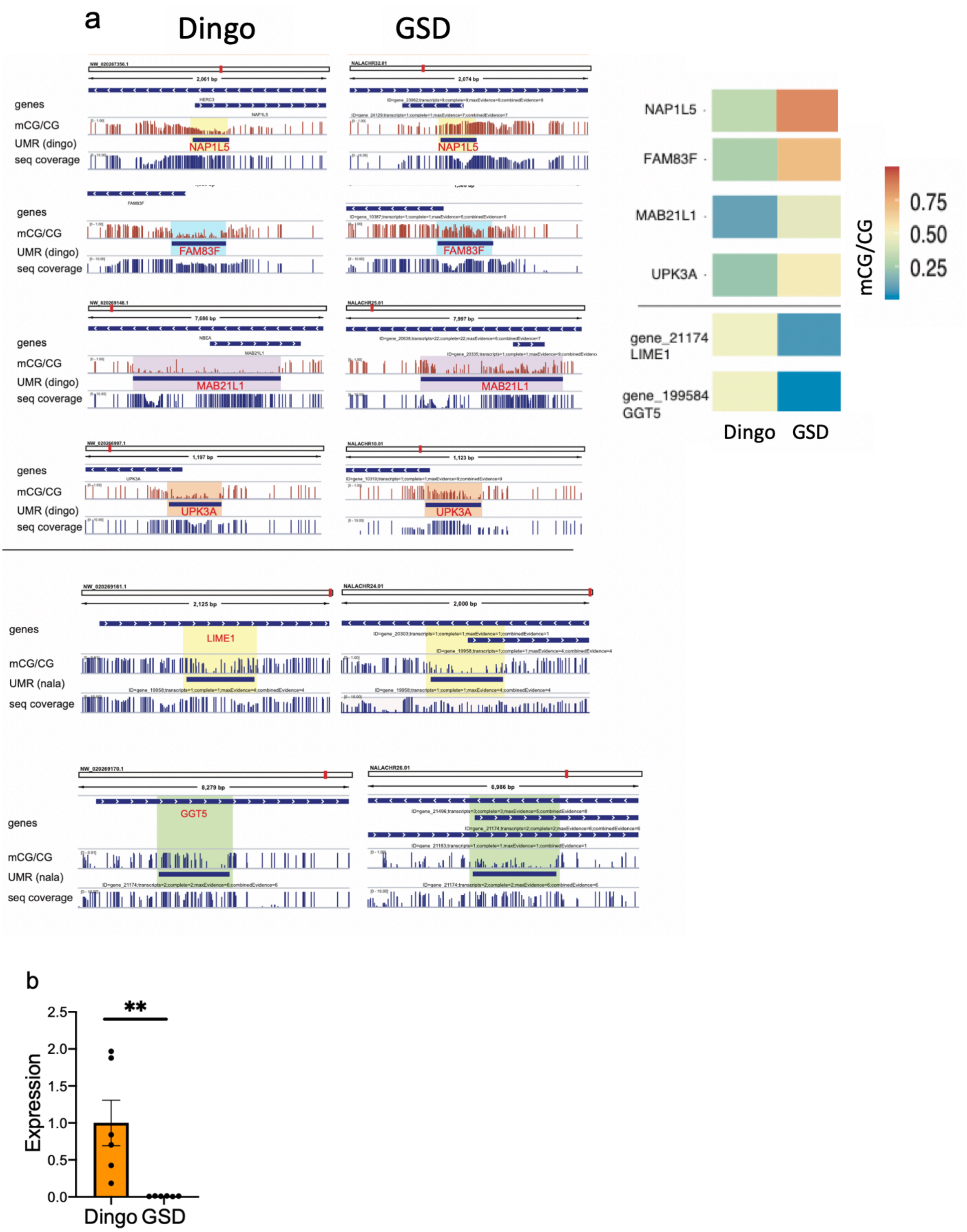


**a,** DNA methylation differences at transcription start sites (TSS) -proximal regulatory regions (UMRs) between the dingo and GSD. Heatmap show DNA methylation levels at TSS-associated UMRs, differentially methylated between dingo and GSD. **b,** Expression difference in *MAB21L1* ($t_{(10)}$ =3.231, P= 0.009). Mean SE is shown on the plot.

**Extended Data Fig. 6: Serum triglyceride and lipase levels between dingo and German Shepherd dog (GSD)**

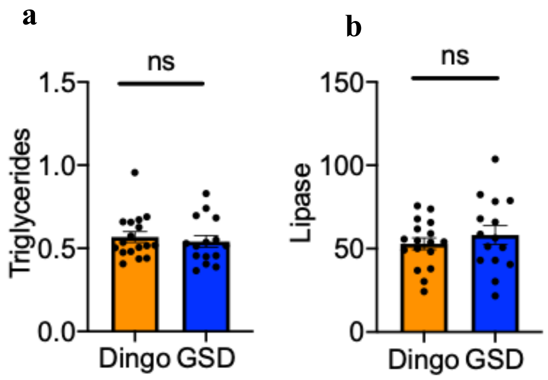


There were no obvious differences in **a,** triglycerides ($t_{30}$ =0.56, P=0.58) and **b,** lipase ($t_{30}$ =0.82, P=0.41). Mean SE is shown on the plot.

#### Extended Data Fig. 7: Microbiome difference between the dingo and German Shepherd Dog (GSD)

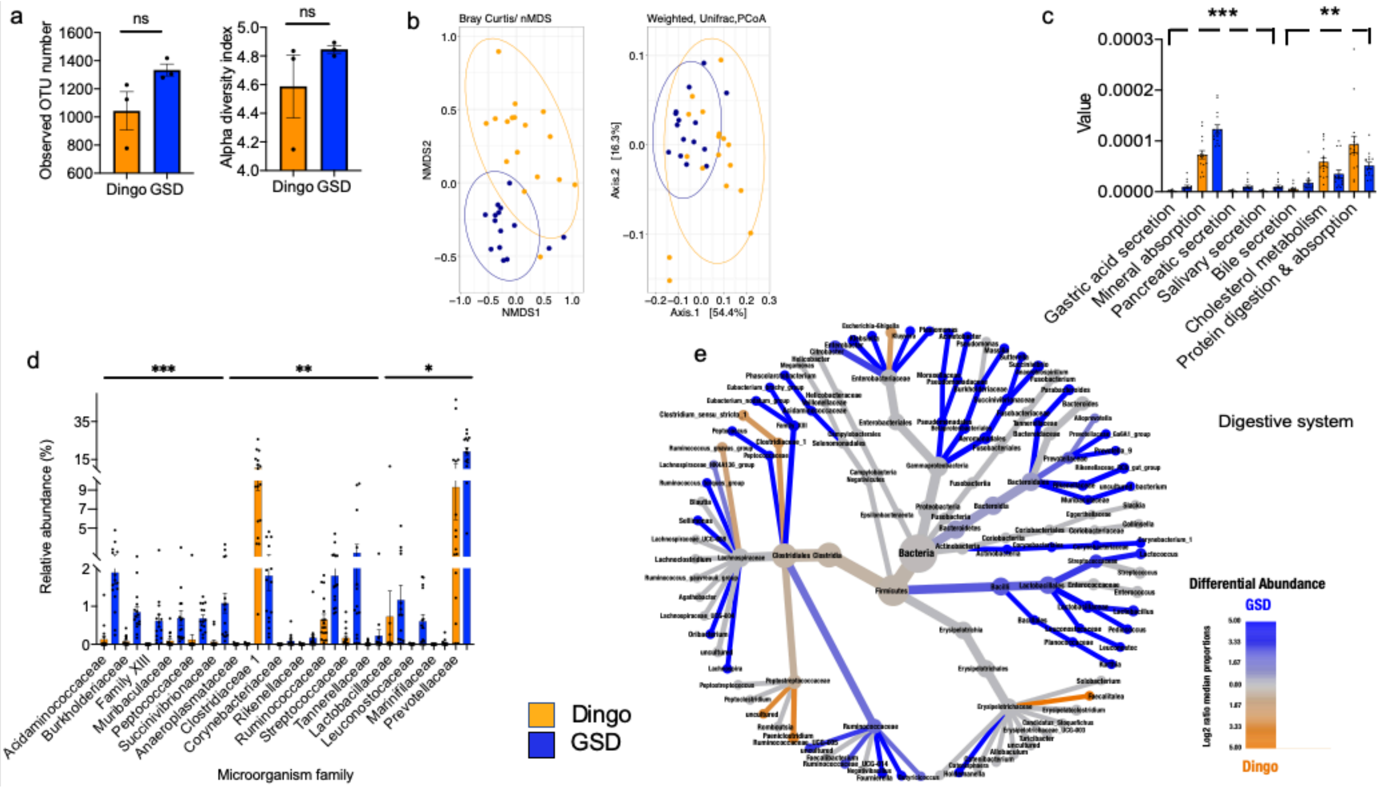


**a**, No significant difference in the microbial richness and diversity at the beginning of the experiment. LHS: Richness, RHS: Diversity**. b,** Microbial communities differ in their community structures at the end of the experiment based on LHS: Bray-Curtis (pairwise.adonis P=0.001) and RHS: weighted Unifrac distances (pairwise.adonis P=0.002). **c**, Metabolic potential of the microbial community predicted using the predictive metagenomic analysis tool, Tax4Fun2. **d**, Relative abundance of significantly different families between Dingo and German Shepherd Dog (GSD) as shown by Analysis of the Composition of the Microbiome (ANCOM) (Benjamini-Hochberg corrected). Mean SE is shown on the barplots. ** shows significance at 0.01, and *** at 0.001. **e**, Differential abundance analysis of top 100 of the most abundant OTUs for both the dingo and GSD, using pairwise comparison between management systems at bloom and harvest. The color of each taxon represents the $\log_2$ ratio of median proportions of relative abundance between the dingo and GSD. Orange taxa are found significantly abundant in the dingo and blue taxa are found significantly abundant in GSD. $\text{Log}_2$ changes are colored, determined using a Wilcoxon rank-sum test followed by a Benjamini–Hochberg (FDR) correction for multiple comparisons. Size of nodes relates to the number of OTUs found within the given taxonomy. Only significant different abundant taxon was visualized by color.
